# Evaluating the Role of Endophyte-Rich Leaves in Protecting Tropical Trees Against a Generalist Herbivore and a Pathogen

**DOI:** 10.1101/2024.04.18.589827

**Authors:** Bolívar Aponte Rolón, A. Elizabeth Arnold, Mareli Sánchez Juliá, Sunshine A. Van Bael

**Affiliations:** Department of Ecology and Evolutionary Biology, Tulane University, 6823 St. Charles Avenue, New Orleans, LA 70118; School of Plant Sciences, Department of Ecology and Evolutionary Biology, Robert L. Gilbertson Mycological Herbarium. Ecosystem Genomics Graduate Interdisciplinary Program, and the Bio5 Institute, University of Arizona, Tucson, AZ 85721; Department of Ecology and Evolutionary Biology and the Bio5 Institute, University of Arizona, Tucson, AZ 85721

**Keywords:** *Atta colombica*, *Calonectria*, foliar fungal endophytes, herbivory, pathogen, symbioses, tropical trees

## Abstract

Plants use chemical compounds and physical barriers to defend themselves against natural enemies. For instance, tough leaves are considered to be better defended than soft leaves and we can measure this spectrum of defenses via leaf functional traits. Leaf traits such as leaf chemistry, lifespan, toughness, and leaf mass per area often are a plant’s first line of defense. Plants with longer lifespans that invest more in leaf tissue and higher leaf mass per area (LMA) typically feature robust constitutive defenses (e.g., leaf toughness, thickness, and dense cell walls) as well. In contrast, plants that invest more in leaf nutrients and have low LMA often invest more in induced defenses. Whether constitutive or induced, leaf traits represent an environmental filter for foliar endophytic fungi (FEF), which may play an additional role in plant defense. Our overarching assumption is that FEF alter leaf fate by interacting directly or indirectly with leaf traits, thus reshaping successive FEF colonization, development of leaf traits and response to plant enemies. To evaluate this hypothesis, we inoculated seedlings of seven tropical tree species that varied in leaf traits with natural and diverse endophyte communities. We confirmed the success of our inoculations of low FEF load (*E-*) and high FEF load (*E+*) treatment groups via culturing and culture-free amplicon sequencing. We then measured leaf removal by leaf-cutter ants and leaf necrosis due to a generalist fungal pathogen. Across the experiment, we observed greater herbivory in the *E-* treatment compared to the *E+* treatment, but no difference in pathogen damage. However, within *E-* and *E+* treatment groups, leaves exposed to pathogen had greater leaf damage than non-exposed leaves. Dimensionality reduction of leaf functional traits (i.e., LMA, toughness, thickness, and anthocyanin levels) revealed relationships among traits and distinct host species characteristics. All leaf functional traits had significant correlations with FEF community composition. In turn, indicator species analyses reveal functional traits and taxonomic identities of FEF associated with high- and low leaf damage by natural enemies, suggesting new insights into cryptic roles of foliar symbionts in extending and modifying plant defenses in tropical forests.

## 3. Introduction

Angiosperms have been successful throughout their evolutionary history by developing strategies against biotic (Feild & Arens, 2005) and abiotic (Leakey & Lau, 2012) selective pressures. Strategies against stressors that damage leaf tissue range from production of secondary metabolites like alkaloid and jasmonic acid (Guerriero et al., 2018; Teoh, 2016) to structures that prevent potential attacks by pests (e.g., thorns/modified leaves)(Hanley et al., 2007). Leaf functional traits such as shape, leaf thickness, leaf strength, leaf mass per area (LMA), and anthocyanins can confer plants with strategies to ward off foliar herbivores and pathogens, which represent key selective evolutionary pressures (J. P. Anderson et al., 2010; Niklas et al., 2023).

Leaf defenses can be placed conceptually along the leaf economic spectrum (LES), which on one end has short-lived leaves with high nitrogen (N) content, low leaf mass per area (LMA), thin leaf blades, and thin cell walls, and on the other end, long-lived leaves with low N content, high LMA, thick leaf blades and thick cell walls (Mason & Donovan, 2015; Wright et al., 2004). Investment in baseline constitutive defenses is associated with longer lifespans (Kitajima et al., 2012; Kitajima et al., 2013), whereas plants that invest little in leaf N content and LMA are potentially able to invest more in induced defense (Kitajima et al., 2013; Poorter & Bongers, 2006; Wright et al., 2004). Such leaf functional traits are expressed differentially across species and are influenced by their life history and the environment they occupy (Kitajima et al., 2013; Wright et al., 2004; Wright et al., 2005). Because they define the chemical, structural, and longevity characteristics of leaves, leaf functional traits also influence associated leaf microbial communities (Saunders et al., 2010; Tellez et al., 2022). Many leaf-associated microbes establish in leaves via horizontal transmission and are thought to alter the physical and chemical traits of leaves (Bael et al., 2017; Chagas et al., 2018; reviewed in Christian et al., 2017). If leaf traits confer selectivity, then plants can gain or lose potential allies in the fight against herbivores and pathogens, ultimately contributing to their ecological and evolutionary success (Friesen et al., 2011).

Leaf microbial communities, such as foliar endophytic fungi (FEF), are found inside the leaf tissue of all lineages of vascular land plants (Currie et al., 2014; Rodriguez et al., 2009). In tropical forests, FEF transmit horizontally through ambient spore fall (Arnold et al., 2000), and newly flushed leaves of such trees are free of FEF. Although FEF generally grow asymptomatically within leaves (Porras-Alfaro & Bayman, 2011), they can modulate leaf functional traits, especially with regard to the expression of secondary metabolites, sensitivity to drought, defense against natural enemies, photosynthetic rates and efficiency (Arnold et al., 2003; Arnold & Engelbrecht, 2007; Bittleston et al., 2011; Estrada et al., 2013; Friesen et al., 2011; Mejía et al., 2014). Such effects have not been examined systematically and quantitatively but are critical to understanding how species interactions in tropical forests relate to plant survival, performance, productivity, and downstream ecosystem services (McGill et al., 2006).

FEF communities occurring in healthy leaves of naturally established tropical trees vary with specific leaf functional traits. Tellez et al. (2022) emphasized tested the hypothesis the abundance, diversity and FEF composition can be explained by leaf traits in the LES and found that FEF were less abundant and diverse in thick, tough long-lived leaves compared to thin, softer leaves from the same forest. The FEF community composition and capacity to produce defensive compounds varied in response to leaf traits in opposing ends of the LES (Tellez et al., 2022). These results extend the LES framework by including diverse and ecologically important fungi. The potential of FEF to alter where plants fall in the LES offers a useful lens to ask: what is the role of FEF in plant defenses against herbivores and pathogens and what trade-offs may be relevant in plant-enemy interactions.

Here we investigated how FEF abundance, diversity and community composition may modulate leaf functional traits and plant’s response to herbivory and pathogen damage. This work builds upon experiments that used single plant species and plant enemies (Estrada et al., 2013; Mejía et al., 2008, 2014) by incorporating seven phylogenetically distinct tropical tree species and two functional classes of plant enemies. We hypothesized that FEF improve leaf defenses against generalist herbivores and pathogens, especially in plants that invest less in constitutive defenses (e.g., thin and short-lived leaves). Alternatively, plants that invest more in constitutive defenses (e.g., thick leaves and long-lived) rely less on FEF improved defenses against plant enemies.

To test our hypothesis we designed an experiment that allowed tropical tree seedlings to be naturally inoculated with FEF. We then measured leaf damage (herbivory and pathogen infection) and a subset of leaf functional traits: leaf mass per area (LMA), leaf thickness (LT), leaf toughness-measured as leaf punch strength (LPS)-, and anthocyanins (ACI), in response to inoculated (high FEF load, *E+*) and non-inoculated (low FEF load, *E-*) treatments. The plant enemies we considered were leaf-cutter ants, *Atta colombica* (Formicidae), a generalist herbivore that, while not consuming the leaves, harvests considerable quantities of leaf tissue to feed underground fungal gardens, and *Calonectria* sp. (Nectriaceae), a generalist foliar pathogen.

We predicted the following: 1) Leaf-cutter ants and *Calonectria* sp. would cause less leaf damage (herbivory through leaf tissue removal and leaf necrosis through pathogen infection, respectively) on leaves with higher FEF abundance, richness and diversity; 2) Tree species with leaf functional traits on the low end of the economic spectrum (e.g., lower leaf mass per area) would have less herbivory and pathogen damage when treated with high FEF loads (*E+*) compared to their low FEF counterparts (*E-*); 3) Conversely, tree species with leaf functional traits of the high side of the economic spectrum (e.g., greater leaf mass per area) treated with high FEF loads (*E+*) would have no differences in herbivory and pathogen damage compared to their low FEF counterparts (*E-*). Lastly, 4) we anticipated that leaves with leaf functional traits on the high end of the economic spectrum would be experience less herbivory by leaf-cutter ants, but low FEF loads (*E-*) in them could increase allurement.

## 4. Materials and Methods

### 4.0.1 Study site and seedling rearing

*Theobroma cacao* (Malvaceae), *Dipteryx* sp. (Fabaceae), *Lacmellea panamensis* (Apocynaceae), *Apeiba membranacea* (Malvaceae), *Heisteria concinna* (Olacaceae), *Chrysophyllum cainito* (Sapotaceae), and *Cordia alliodora* (Cordiaceae) were chosen due to their variation in leaf functional traits (J.Wright *unpublished data*). All occur naturally at Barro Colorado Island (BCI) in central Panama (9°050N, 79°450W), where we collected seeds from the forest floor from multiple maternal sources in January - April 2019. Average annual precipitation at BCI is 2,600 mm and the pronounced wet season ranges from May to December (Leigh et al., 1996). In preparation for the experiment, seeds were surface sterilized by soaking in water and rinsing in sodium hypochlorite (NaClO) and ethanol (EtOH). Seeds from each species had a species-specific sterilization protocol due to the variation in sizes and seed coats (see Supplementary Materials). Seed germination and the subsequent experiment were carried out at the Santa Cruz Field Facility of the Smithsonian Tropical Research Institute in Gamboa, Panama (9°070N, 79°420W). We germinated and reared seedlings in a clean and shaded greenhouse where we enclosed four tables with a PVC pipe frame and covered them with a 3 mil clear plastic sheet, for a total of two plastic enclosures with two tables each. The enclosures allowed us to grow plants at ambient temperature and natural light while providing protection from rain and most fungal spores, thus yielding zero to low FEF densities n plants that were not actively inoculated through our manipulations(see below) (Bittleston et al., 2011). We cleaned table surfaces and walls of the enclosures on a weekly basis with 70% EtOH and 0.5 % NaClO. We germinated seedlings in sterilized trays containing a 3:1 mix of soil and river sand that was autoclaved for two one-hour cycles at 121°C prior to planting. Individual seedlings were transferred from germination trays to a 24-cell tray (each cell ∼380 mL) containing the same autoclaved soil and sand mixture. We took precautions to extract complete root systems from the seedlings. See Supplementary Materials for further details on plastic tray and pot sterilizations protocols.

Seedlings reached a minimum of 5-6 true leaves before endophyte inoculation. We placed seedlings on separate tables designated for spore fall inoculation and non-inoculated treatment groups within the enclosures. Seedlings of the same species but different treatment groups were in the same enclosure. We watered seedlings at the soil level to minimize endophyte spore germination in the enclosures (Arnold et al., 2003).

### 4.0.2 Fungal endophyte inoculation

To inoculate seedlings with FEF, we took 10 individual seedlings of each species and exposed them over 10 nights to natural spore fall in the forest understory to achieve a high FEF load (*E+*) and 10 plants were kept inside the greenhouse enclosure to maintain a low FEF load (*E-*). Plants exposed to spore fall were placed on a table near (∼10 m) the forest edge at dusk (∼18:00 hours) and returned to the greenhouse at dawn (∼07:00 hours) (Bittleston et al., 2011). We sprayed the *E+* seedlings with water to simulate rain and to promote endophyte spore germination and infection of leaves. Low FEF plants (*E-*) were watered only at the soil level and shuffled and moved inside the greenhouse to simulate similar treatment as *E+* seedlings, but without spore fall exposure.

### 4.0.3 Leaf trait measurements

Three mature leaves were haphazardly collected 7 - 10 days after fungal inoculation from individuals in each treatment (*E+*, *E-*) with 5 - 6 true leaves. Anthocyanin (ACI) content and leaf thickness (LT) were measured while the leaf was still attached to the plant. We measured anthocyanin content with an ACM-200plus anthocyanin content meter (Opti-Sciences Inc. Hudson, New Hampshire, U.S.A.) on three haphazardly selected locations (working from the petiole out to the leaf tip) on the leaf surface of three haphazardly selected leaves for a total of nine measurements per plant (Tellez et al., 2022). To account for leaf thickness, the ACM-200 calculates an anthocyanin content index (ACI) value from the ratio of % transmittance at 931 nm/% transmittance at 525 nm (Tellez et al., 2016). On compound leaves (i.e., *Dipteryx* sp.) we measured at three different leaflets. Leaf thickness (μm) was measured with a Mitutoyo 7327 Micrometer Gauge (Mitutoyo, Takatsu-ku, Kawasaki, Japan) at six different points on the leaf lamina; at the base, mid-leaf and tip on both sides of the mid-vein, taking care to avoid major and secondary veins. After ACI and leaf LT measurements were completed, we removed the leaves from their stems, placed them inside a zip-top plastic bag in an ice chest and moved them to the lab for further measurements. Leaf punch strength (LPS) was measured with an Imada DST-11a digital force gauge (Imada Inc., Northbrook, IL, United States) by conducting punch-and-die tests with a sharp-edged cylindrical steel punch (2.0 mm diameter) and a steel die with a sharp-edged aperture of small clearance (0.05 mm). Leaf punch measurements were taken at six locations on each leaf by puncturing the lamina at the base, mid-leaf and tip on both sides of the mid-vein, and avoiding minor leaf veins when possible (Tellez et al., 2022). Once LPS was measured, we used a 7 mm diameter hole punch to obtain three leaf disks per leaf for leaf mass per area (LMA) (see Supplementary material for details). Disk punches dried at 60 °C for 48-72 hours before being weighed for dry mass immediately.

### 4.0.4 Leaf tissue preparation for molecular work

The same three leaves were pooled and used to profile FEF abundance, richness, and community composition via amplicon sequencing (Illumina MiSeq). The leaf tissue remaining after the leaf trait measurements had the main vein and margins excised so that only the laminae remained. The laminae were haphazardly cut into 2 x 2 mm segments, enough to obtain a total of 16, and surface sterilized by sequential rinsing in 95% ethanol (10 s), 0.5 NaOCl (2 mins) and 70% ethanol (2 mins) (Arnold et al., 2003; Higgins et al., 2014; Tellez et al., 2022). Leaves were then surface-dried briefly under sterile conditions. Sixteen leaf segments per leaf (i.e., forty-eight leaf segments per plant) were plated on the surface of 2% malt extract agar (MEA) in Petri dishes (60 mm), sealed with Parafilm M (Bemis Company Inc., U.S.A.) and incubated at room temperature. The cultured leaf segments were used to estimate FEF colonization of *E-* and *E+* leaves. The presence or absence of endophytic fungi in culture was assessed 7 days after plating. The remaining sterilized laminae were preserved in sterile 15 mL tubes with ∼ 10 mL CTAB (1 M Tris–HCl pH 8, 5 M NaCl, 0.5 M EDTA, and 20 g CTAB). Leaf tissue in CTAB was used for amplicon sequencing (described in detail below). All leaf tissue handling was performed in a biosafety cabinet with all surfaces sterilized by exposure to UV light for 30 minutes and cleaned sequentially in between samples with 95% ethanol, 0.5% NaOCl and 70% ethanol to prevent cross contamination. Leaf tissue in CTAB was stored for 2 months at room temperature prior to storage at -80 °C for 3 months preceding DNA extraction

### 4.1 Amplicon sequencing

In preparation for DNA extraction, we decontaminated all instruments, materials, and surfaces with DNAway (Molecular BioProducts Inc., San Diego, CA, United States), 95% ethanol, 0.5% NaOCl, and 70 % ethanol, and subsequently treated with UV light for 30 minutes in biosafety cabinet. We used sterile equipment and pipettes with aerosol-resistant tips with filters in all steps before amplification. From each sample in CTAB in we transferred 0.2 – 0.3 g of leaf tissue into duplicate sterile 2mL tubes, resulting in 2 subsamples. Total genomic DNA from each subsample was extracted as described in U’Ren & Arnold (2017). In brief, we added two sterile 3.2 mm stainless steel beads to each tube and proceeded to lyophilize samples for 72 hours to fully remove CTAB from tissue. Then we submerged the sample tubes in liquid nitrogen for 30s and homogenized samples to a fine powder for 45 s in FastPrep-24 Tissue and Cell Homogenizer (MP Biomedicals, Solon, OH, USA). We then repeated the decontamination procedure described before and used the QIAGEN DNeasy 96 PowerPlant Pro-HTP Kit (U’Ren & Arnold, 2017) (QIAGEN, Valencia, CA, USA) to extract total genomic DNA. We pooled the subsamples for each individual sample before amplification. We followed a two-step amplification approach previously described by Sarmiento et al. (2017) and ÚRen & Arnold (2017). We used a separate set of sterile pipettes, tips, and equipment to reduce contamination. We used a designated PCR area to restrict contact with pre-PCR materials (Oita et al., 2021). Used primers for the fungal ITSrDNA region, ITS1f (5’-CTTGGTCATTTAGAGGAAGTAA-3’) and ITS4 (5’-TCCTCCGCTTATTGATATGC-3’) with modified universal consensus sequences CS1 and CS2 and 0–5 bp for phase-shifting. Every sample was amplified in two parallel reactions containing 1-2 µL of DNA template (U’Ren & Arnold, 2017; see also Tellez et al., 2022). We visualized PCR (PCR1) reactions with SYBR Green 1 (Thermo Fisher Scientific, Waltham, MA, USA.) on a 2% agarose gel (Oita et al., 2021). Based on band intensity, we combined parallel PCR1 reactions and diluted 5 µL of amplicon product with molecular grade water to standardize to a concentration of 1:15 (Sarmiento et al., 2017; Tellez et al., 2022). We included DNA extraction blanks and PCR1 negatives in this step. We used 1 µL of PCR1 product from samples and negative control for a second PCR (PCR2) with barcode adapters (IBEST Genomics Resource Core, Moscow, ID, USA). Each PCR2 reaction (total 15 µL) contained 1X Phusion Flash High Fidelity PCR Master Mix, 0.075 µM of barcoded primers (forward and reverse pooled at a concentration of 2 µM) and 0.24mg/mL of BSA following Sarmiento (2017) and U’Ren & Arnold (2017). Before final pooling for sequencing, we purified the amplicons using Agencourt AMPure XP Beads (Beckman Coulter Inc, Brea, CA USA) to a ratio of 1:1 following the manufacturer’s instructions. The products were evaluated with Bio Analyzer 2100 (Agilent Technologies, Santa Clara, CA, USA) (Tellez et al., 2022). We quantified the samples through the University of Arizona Genetics Core, and subsequently diluted them to the same concentration to prevent over representation of samples with higher concentration (Sarmiento et al., 2017). Amplicons were normalized to 1 ng/µL, then pooled 2 µL of each for sequencing. No contamination was detected visually or by fluorometric analysis. To provide robust controls we combined 5 µL of each PCR1 negative and the DNA extraction blanks and sequenced them as samples. Ultimately, we combined samples into a single tube with 20 ng/µL of amplified DNA with barcoded adapters for sequencing on the Illumina MiSeq platform with Reagent Kit v3 (2 × 300 bp) following protocols from the IBEST Genomics Resource Core at the University of Idaho, USA. Again, we included the DNA extraction blanks and two PCR1 negatives and sequenced with samples.

#### 4.1.1 Mock Communities

We processed and sequenced a mock community following the methods described above. We had two aims: to understand the relationship between read abundance and biological abundance, and to determine whether primer bias might exclude fungal lineages of interest from our sequence data. We used two mock communities that consisted of PCR product from DNA extractions of 32 phylogenetically distinct fungi, representing lineages that are typically observed as endophytes: Ascomycota, Basidiomycota, fungi traditionally classified as Zygomycota, and Chytridiomycota (Oita et al., 2021; see Daru et al., 2019 for details). In brief, we sequenced six replicates of the mock community with equimolar concentrations of DNA from all 32 fungal taxa, and another six replicates of the mock communities with tiered concentrations of DNA from the same fungal taxa (see Daru et al., 2019). Read abundance from tiered communities was positively associated with the expected read number (*R^2^_adj_*= 0.87, *p* < .0001, see Fig. S9), and all major fungal lineages present in the mock community were detected (data not shown). Henceforth, we used read abundance as a relevant proxy for biological OTU abundance (U’Ren et al., 2019).

#### 4.1.2 Bioinformatic analyses

We used VSEARCH (v2.14.1) for *de novo* chimera detection, dereplication, and assignment of sequences to operational taxonomic units (OTU). VSEARCH is an open-source alternative to USEARCH that uses an optimal global aligner (full dynamic programming Needleman-Wunsch), resulting in more accurate alignments and sensitivity (Rognes et al., 2016). For mock communities and experimental samples, we used forward reads (ITS1) for downstream bioinformatics analyses due to their high quality, rather than reverse reads (ITS4). Following Sarmiento et al. (2017), we concatenated all reads in a single file and used FastQC reports to assess Phred scores above 30 and determine the adequate length of truncation. We processed 892,713 sequence reads from mock communities and 3,778,081 from experimental samples. We truncated mock community and experimental sample reads to a length of 250 bp with command `fast_trunclen` and filtered them at a maximum expected error of 1.0 with command `fast_maxee`. We then clustered unique sequence zero radius OTUs (that is, zOTUs; analogous to amplicon sequence variants (Callahan et al., 2016)), by using commands `derep_fulllength` and `minseqlength` set at 2. Sequentially we denoised and removed chimeras from read sequences with commands `cluster_unoise`, and `uchime3_denovo`, respectively. Finally, we clustered zOTUs at a 95% sequence similarity with command `usearch_global` and option `id` set at 0.95. At that point, 3,035,960 sequence reads from experimental samples remained. Taxonomy was assigned with the Tree-Based Alignment Selector Toolkit [v2.2; Carbone et al. (2019)] by placing unknowns within the Pezizomycotina v2 reference tree (Carbone et al., 2017), and blasting against the UNITE database via the ribosomal database project (RDP) classifier. A total of 2147 OTUs were obtained, with the combined taxonomic data sets revealing 68.6% Ascomycota, 26.8% Basidiomycota, and other fungal lineages either rare (e.g., <0.05% Chytridiomycota, Mortierellomycota) or unidentified (4.2 %). Only OTUs representing Ascomycota were used for downstream statistical analyses since foliar endophyte communities in tropical trees are dominated by Ascomycota (Arnold & Engelbrecht, 2007). For each OTU identified, we removed laboratory contaminants from experimental samples by subtracting the average read count found in control samples from the DNA extraction and PCR steps. Our analysis of mock communities allowed use to identify and remove false OTUs from experimental samples, those with fewer than 10 reads, leading us to 0.1% of the read relative abundance across all samples (Oita et al., 2021). Three experimental samples from *Theobroma cacao* (*n*=2) and *Apeiba membranacea* (*n*=1) were removed from all analyses due to incomplete entries. After pruning OTU with zero reads from experimental samples, we identified 260 OTUs found exclusively in control (*E-*) plants (*n*=78) and deemed them as artifacts resulting from greenhouse conditions. They were eliminated from treatment (*E+*) plants across all species. We converted reads for each fungal OTU to proportions of total sequence abundance per sample to reduce differences in sampling effort, following previous studies (Weiss et al. (2017)). We then removed singletons and obtained an average of 2,464,558 sequence reads in 529 Ascomycota OTUs across 156 experimental samples of 7 tree species. All analyses after taxonomic assignment were performed in R [v. 4.3.3; R Core Team (2024)] using the `phyloseq` package (McMurdie & Holmes, 2013) and custom scripts (see Supplementary Material).

#### 4.1.3 Herbivore assays

To assess leaf-cutter ant damage, we collected one extra leaf per plant per treatment, at the same time we collected samples for leaf trait measurements, and introduced it to an actively foraging leaf-cutter ant colony for a two-hour assay. We presented leaf-cutter ant colonies with a choice of an *E+* or an *E-* leaf on one disposable plastic plate next to an active nest trail. Carefully, we collected and placed debris from the trail leading up to the plate to encourage foraging in the plate. We initiated the ant assay as soon as one ant entered the plate and explored the leaf contents for at least 10-20 seconds. Every five minutes we took a digital photo of the choice arena until about 75% of the leaf content of one of the leaves was taken. We used the digital photo at time zero and at the end of trial to quantify the leaf area removed using ImageJ [v1.52r; Schneider et al. (2012)]. Ant recruitment was estimated by counting individuals in the choice arena throughout trial event.

#### 4.1.4 Pathogen assays

For the pathogen assays, we introduced an agar plug inoculated with hyphae of *Calonectria* sp. (*P+* treatment), and an agar plug without the pathogen (*P-* control) to similarly aged/sized leaves that were still on plants (i.e., were not harvested) within 10-14 days after endophyte inoculations (Gilbert & Webb, 2007). Leaves with the *P+* or *P-* treatment were misted with sterile water two times a day (morning and afternoon) to maintain moisture. After four days, we removed the plugs and took digital photos to analyze leaf area damage using ImageJ [v1.52r; Schneider et al. (2012)].

#### 4.1.5 Replication Statement

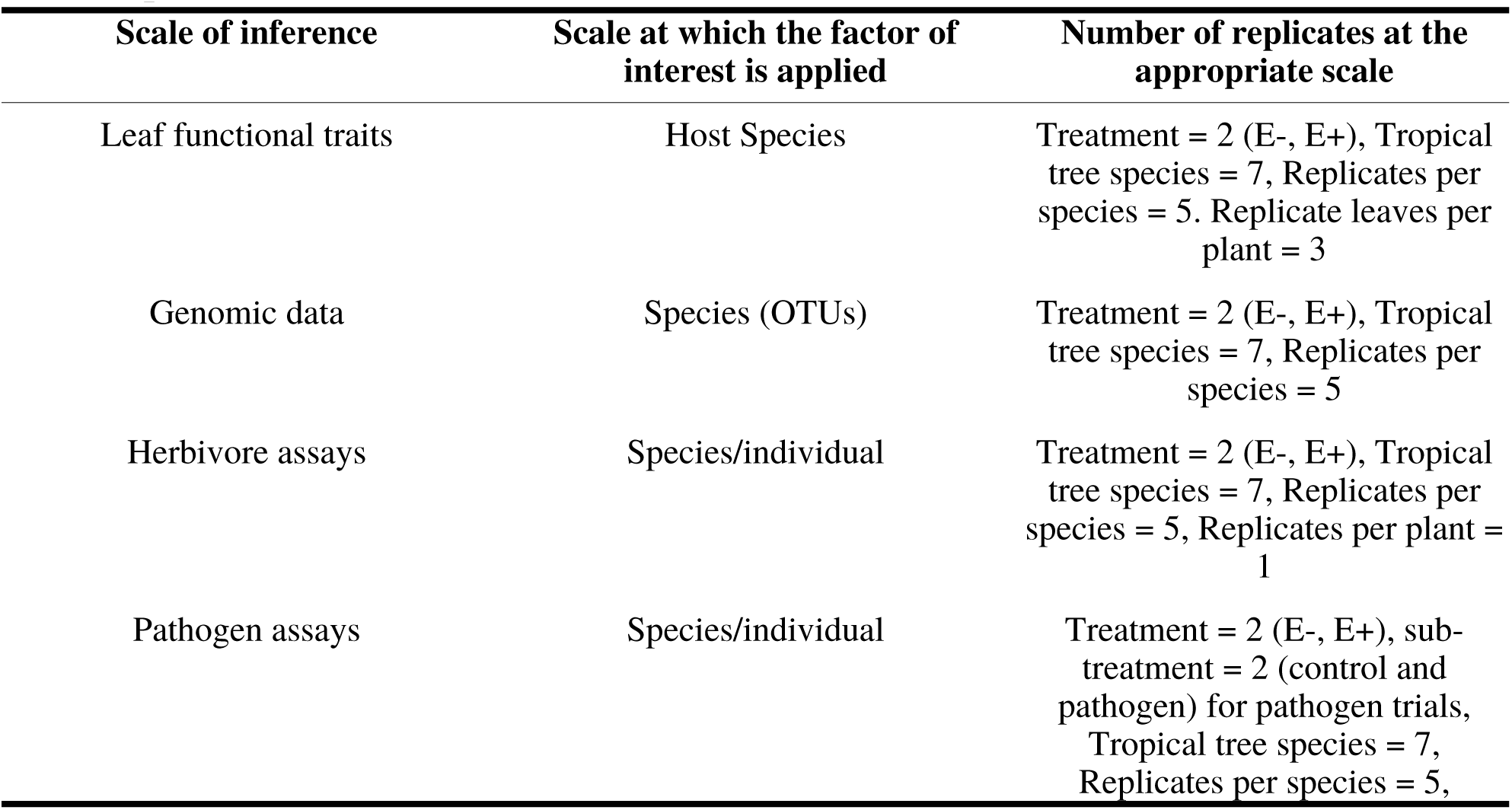
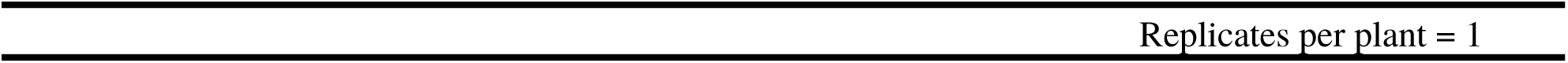

| Scale of inference | Scale at which the factor of interest is applied | Number of replicates at the appropriate scale |
| --- | --- | --- |
| Leaf functional traits | Host Species | Treatment = 2 (E-, E+), Tropical tree species = 7, Replicates per species = 5. Replicate leaves per plant = 3 |
| Genomic data | Species (OTUs) | Treatment = 2 (E-, E+), Tropical tree species = 7, Replicates per species = 5 |
| Herbivore assays | Species/individual | Treatment = 2 (E-, E+), Tropical tree species = 7, Replicates per species = 5, Replicates per plant = 1 |
| Pathogen assays | Species/individual | Treatment = 2 (E-, E+), sub-treatment = 2 (control and pathogen) for pathogen trials, Tropical tree species = 7, Replicates per species = 5, |

#### 4.1.6 Statistical Analyses

We explored how leaf functional traits and FEF correlated to herbivory and pathogen damage on leaves. We present the analyses for each tree species at the leaf and at the plant level. Leaf functional traits are presented at the leaf level, while FEF data are presented at the host species level (plant), consistent with our whole-plant inoculation approach. In analyses where leaf functional traits and FEF are combined we used averages of leaf functional traits.

First, we compared the means of herbivory (%) damage, and leaf functional traits for each species and treatment groups using paired two-sided Student’s t-Test and analysis of variance (ANOVA) with the `compare_means` and `stat_compare_means` functions from the `ggpubr` package in R (Kassambara, 2023b), which wrap and extend the `anova` and `t.test` functions from the `stats` package (R Core Team, 2024) and facilitate plotting. Additionally, to facilitate comparison between group levels we used the `pairwise_t_test` function from the `rstatix` package (Kassambara, 2023a) when we compared pathogen (%) damage means. This function also wraps and extends base *R* functions. We adjusted *p* values to account for false discovery rates in multiple comparisons by using “BH” method (Benjamini & Hochberg, 1995).

Secondly, we calculated a Bray-Curtis dissimilarity matrix with our OTU relative abundance data and computed a distance-based redundancy analysis (dbRDA) by applying the `dbRDA` function in the `vegan` package to our dissimilarity matrix (Oksanen et al., 2022). We computed forward model selection for dbRDA analysis with the `ordistep` function which selects terms based on *p* values (Blanchet et al., 2008; Oksanen et al., 2022). We started with our initial model containing only the intercept (dissimilarity_matrix ∼ 1) and setting the functions arguments to the following: `scope` = formula(*m*), where *m* is the formula with a defined range including leaf functional traits, tree species and treatment groups; `Pin` = 0.5, `Pout` = 0.05, `trace` = T, `permutations` = how(nperm = 999), `steps` = 50.The dbRDA is considered analogous to a permutational analysis of variance (PERMANOVA) with non-Euclidean distance (M. J. Anderson, 2017; McArdle & Anderson, 2001). Its corresponding visualizations appropriately illustrate underlying patterns of compositional differences (M. J. Anderson, 2017; Legendre & Anderson, 1999; McArdle & Anderson, 2001). We used the `anova.cca` function to assess the marginal significance of constraining variables (Legendre et al., 2011; Legendre & Legendre, 2012; Oksanen et al., 2022), applied here to reveal associations between leaf functional traits and FEF communities in host tree species and treatment groups. To corroborate homogeneous dispersion of variances of host species and treatment groups we used a permutational analysis of multivariate dispersion (PERMDISP) using the `betadisper` with parameter `type = “median”`, and `permutest` functions from `vegan` with with 999 permutations (Oksanen et al., 2022). We compared differences in the dispersion of the FEF communities among species and treatments with post-hoc Tukey’s tests.

Thirdly, we arbitrarily designated percent leaf damage in herbivore assays as high (>70%), medium (31-69%) and low (<30%) and in pathogen assays as high (>30%) and low (<30%). These categories allowed us to explore correlations between host tree species and specific FEF OTUs. To achieve this we used the `multipatt` function from the `indicspecies` package in *R* (De Cáceres & Legendre, 2009). We calculated the point biserial correlation coefficient for each OTU at all tree species and treatment group combinations by applying the `multipatt` function with arguments `func`= “r.g” and `control` = how(nperm=999) to our OTU abundance matrix (De Cáceres & Legendre, 2009). Like with other statistical tests performed, we adjusted *p* values to account for false discovery rates in multiple comparisons by using “BH” method (Benjamini & Hochberg, 1995) in the `p.adjust` function from the `stats` package (R Core Team, 2024). We then filtered the adjusted *p* value with a cutoff of < .05.

We used Principal Component Analysis (PCA) to reduce dimensions among covariates and reveal underlying interactions between covariates that could influence herbivory and pathogen damage. The PCA was computed using the `prcomp` function in R statistical software (R Core Team, 2024). A complete PCA was computed with variables ACI, LT, LPS, and LMA. We then proceeded to compute a PCA with the data from leaves of plants used in the herbivory (*n* = 210) and pathogen assays (*n* = 192). We then took from the herbivory and pathogen PCA the principal components that explained the most variance (PC1 and PC2) and regressed them to herbivory (%) and pathogen damage (%). We also regressed ACI, LT, LPS, LMA and Shannon diversity index to logit transformed herbivory (%) and pathogen damage (%). We used the `logit` function from the `car` package for logit transformation of variables and the `lm` function from the `stat` package for simple linear regressions (Fox & Weisberg, 2019; R Core Team, 2024).

Lastly, to test how leaf functional traits and FEF communities interact to influence herbivory and pathogen damage in tropical tree species, we used a generalized linear mixed model (GLMM) with herbivory and pathogen damage percentage (logit transformed) as the response variable. To determine which fixed effects to include in the GLMMs we used the `vif` function from the `car` package in *R* to calculate the variance inflation factor for all explanatory variables (ACI, LT, LPS, LMA, and Shannon diversity index) (Fox & Weisberg, 2019; R Core Team, 2024).

Complementary to this, we calculated Pearson’s coefficient for each pair of leaf functional traits with by creating a correlation matrix and applying the `cor` function from the `stats` package to assess correlations among traits (R Core Team, 2024). We opted to maintain explanatory variables LT and LMA, and exclude ACI and LPS from subsequent general linear models (GLMMs) due to high collinearity between the two variables with LMA, *r*(1112) = 0.68, *p* < .0001 and *r*(1113) = 0.65, *p* < .0001, respectively. We modeled only main effects with explanatory variables, we did not model interactions effects to avoid overfitting models. We used a restricted maximum likelihood estimates for model fit with the `lme` function from the `nlme` package (J. Pinheiro et al., 2023; J. C. Pinheiro & Bates, 2000). For our logit herbivory GLMMs we used tree species as a random effect and modeled tree species variance structure with the `varIdent` argument (J. Pinheiro et al., 2023; J. C. Pinheiro & Bates, 2000). For our logit pathogen damage GLMMs we use tree species as a random effect and modeled a nested variance structure for pathogen treatment within treatment groups per species with the `varIdent` argument (J. Pinheiro et al., 2023; J. C. Pinheiro & Bates, 2000). We manually compared and selected models based on Akaike Information Criterion (AIC) with a penalty of 2 degrees of freedom (ΔAIC) with the `model.sel` function from the `MuMIn` package (Bartoń, 2023). We selected the best-fit model based on the lowest value obtained.

## 5. Results

Inoculation of seedlings was successful. Seedlings exposed to forest spore fall (i.e., *E+*) had a significantly higher proportion of leaf segments colonized by FEF across all species (data from cultures, Fig. S1). Similarly, molecular data set showed that seedlings with *E+* treatment had a significantly higher FEF relative abundance (paired, two-sided *t*-tests, *p* < .05) for all tree species when compared to the *E-* treatment (Fig. 1a). Despite these significant differences, there was a high degree of variability in FEF relative abundance within each treatment type (Fig. 1).

**Figure 1.**
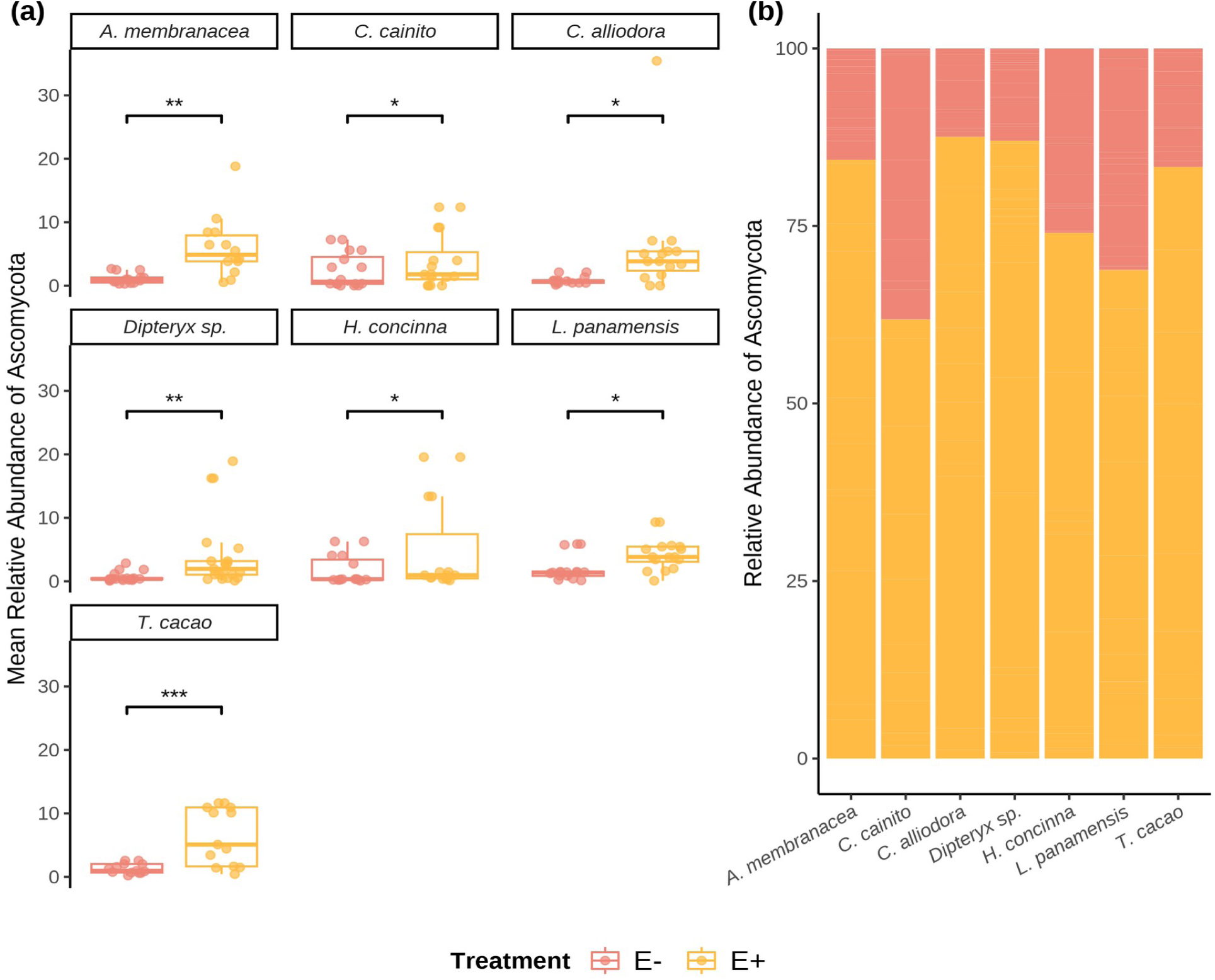
Relative abundance (RA) of Ascomycota OTUs of seven tree species used in the study. (a) Box plots show individuals’ RA and its distribution by species. (b) The RA of OTU’s by treatment withing each tree species. Pink outlined box plots represent low endophyte (*E-*) treatment and yellow filled represent high endophyte (*E+*) treatment. Relative abundance is the percentage of endophyte colonization within individuals of the same species. Statistical significance was calculated using a two-sided Student’s t-Test. Significance levels are represented by *ns* (not significant) and asterisks [*p* < .05 (*), *p* < .01 (\**\*),* p < .001 (***)*, p <* .001 (***), and *p* < .0001 (****)].

We observed general differences in leaf functional traits among species (Table 1). Anthocyanins (ACI) and leaf punch strength (LPS) did not differ between treatments (*E-* and *E+*) when we combined all host species (Fig. S2a and Fig. S4a respectively). For LT and LMA we saw significant differences between *E-* and *E+* treatment groups when we combined all host species (Fig S3a and Fig. S5a, respectively). As predicted, we did observe lower herbivory in the *E+* treatment compared to the *E-* treatment when we combined all host species, *t*(34) = 2.23, *p* = 0.033 (Fig. 2a). We did not observe differences in pathogen damage between *E-* and *E+* treatments when we combined all host species (Fig. 2b). However, within the *E-* treatment group, leaves exposed to *Calonectria* sp. suffered greater damage (*M* = 17.37, *SD* = 12.57) compared to non-exposed leaves (*M* = 3.49, *SD* = 5.42), *t*(31) = -7.19, *p* < .0001 (Fig. 2b). The same pattern arose for the *E+* treatment group: exposed leaves (*M* = 20.04, *SD* = 15.72), non-exposed leaves (*M* = 7.17, *SD* = 17.48), *t*(31) = -3.26, *p* < .01 (Fig. 2b). At the species level, herbivory and pathogen damage did not differ between treatment groups (Fig. S6a-S6b).

**Figure 2.**
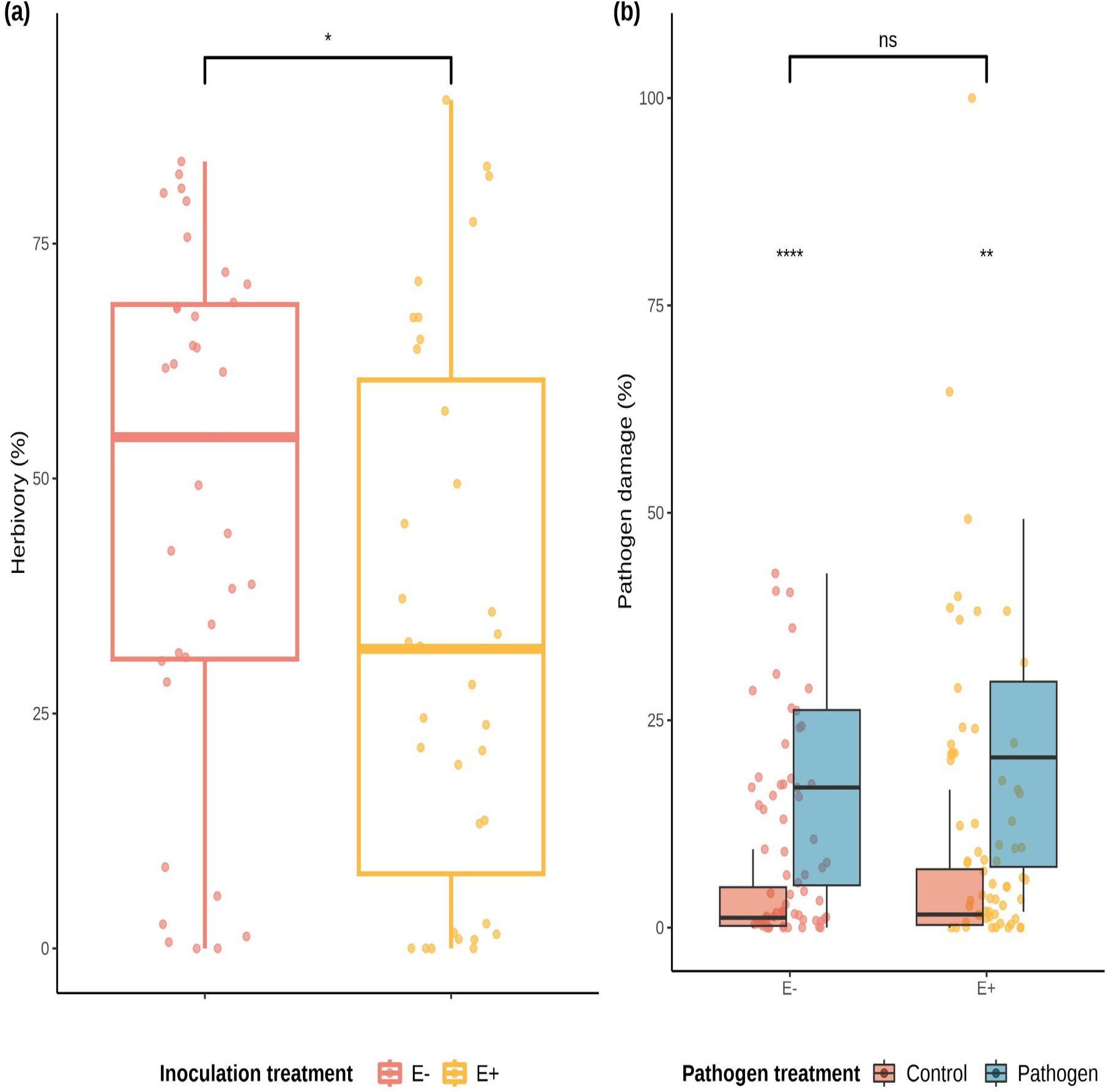
Distributions of values and means of herbivory (%) damage caused by *Atta colombica* in treatment groups (*E-* and *E+*) and tree species. a) Comparison of herbivory (%) means between treatment groups across individuals of all species. outlined box plots represent low FEF group (*E-*) and yellow outlined box plots represent high FEF group (*E+*). b) Comparison of pathogen (%) means between treatment groups across individuals of all species. Maroon filled violins represent control leaves and blue filled violins represent pathogen treated leaves. Statistical significance was calculated using a two-sided Student’s t-Test. Significance levels are represented by *ns* (not significant) and asterisks [*p* < .05 (*), *p* < 0.01 (**), p < .001 (***)*, p =* .001 (***), and *p* < .0001 (****)].

**Table 1:**
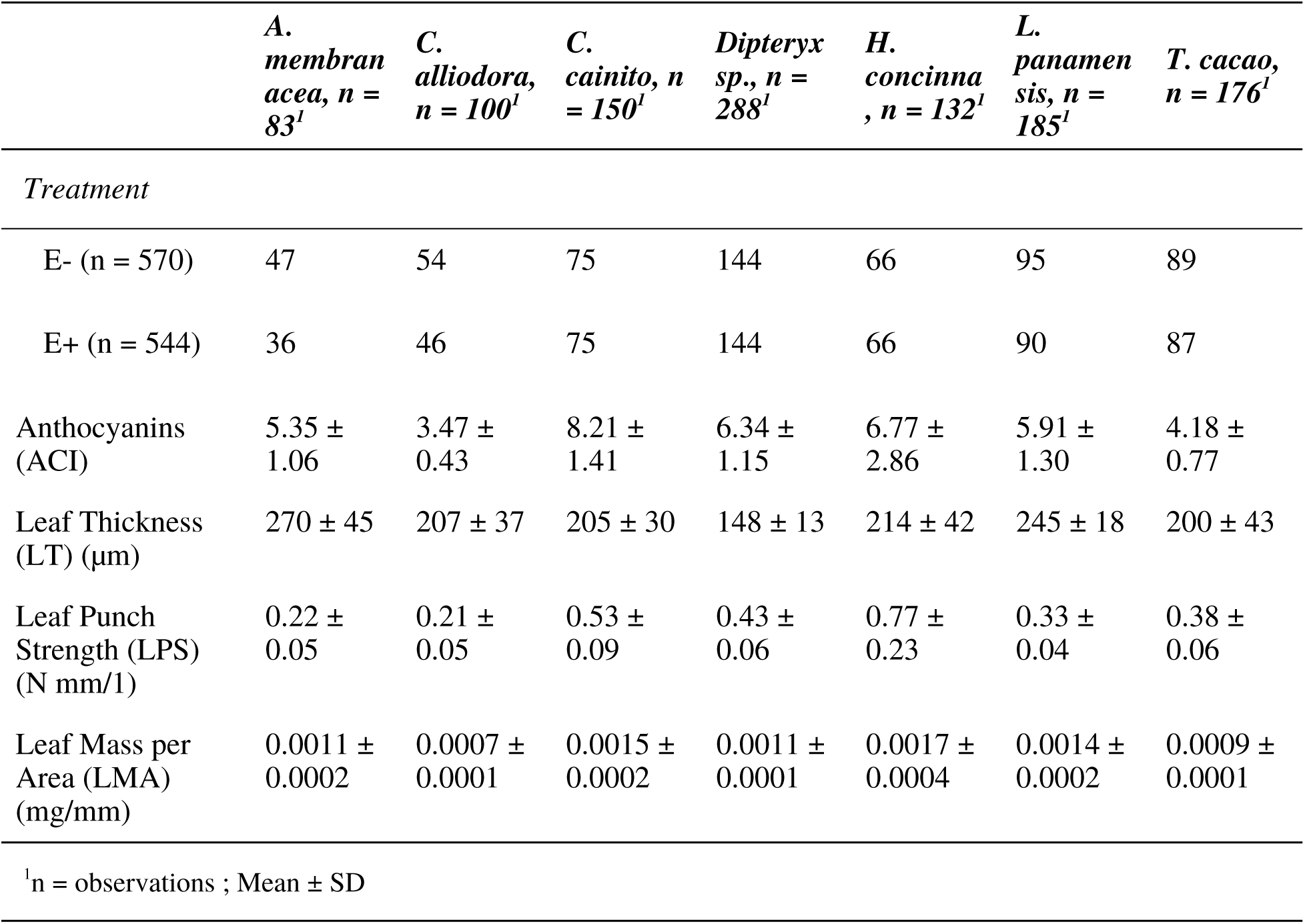
Summary statistics for the leaf functional traits.

All leaf functional traits are significantly correlated with FEF community composition (Fig.3). We used dbRDA to understand FEF community composition across host species and treatment groups. The analyses revealed that 6.34% of the overall variance in FEF communities was constrained for by the leaf functional traits. The first axis (dbRDA1) explained 49% and the second axis (dbRDA2) explained 21.3% of the constrained variance (Fig. 3). We observed a high degree of overlap between FEF communities (Fig. 3), indicating that the communities are similar in composition across host species and treatment groups, at least at the degree of resolution provided by ITS data (Fig. 3). Nonetheless, we also observed tight clustering of FEF communities in *C. cainito* and *L. panamensis*, emphasizing a distinct composition of FEF OTUs within the subspace of FEF identified. Other host species showed greater variation in FEF composition. Our PERMDISP analyses showed no significant differences in host species group dispersion (*F*∼6, 149∼ = 0.717, *p* < .63), observed differences are were to location. Unsurprisingly, the dispersion for *E-* and *E+* treatment groups was significantly different (*F*∼1, 154∼ = 5.09, *p* = .03). The post Tukey tests revealed dispersion of host species were not significantly different at the α < .05 level, while treatment group dispersions were.

**Figure 3.**
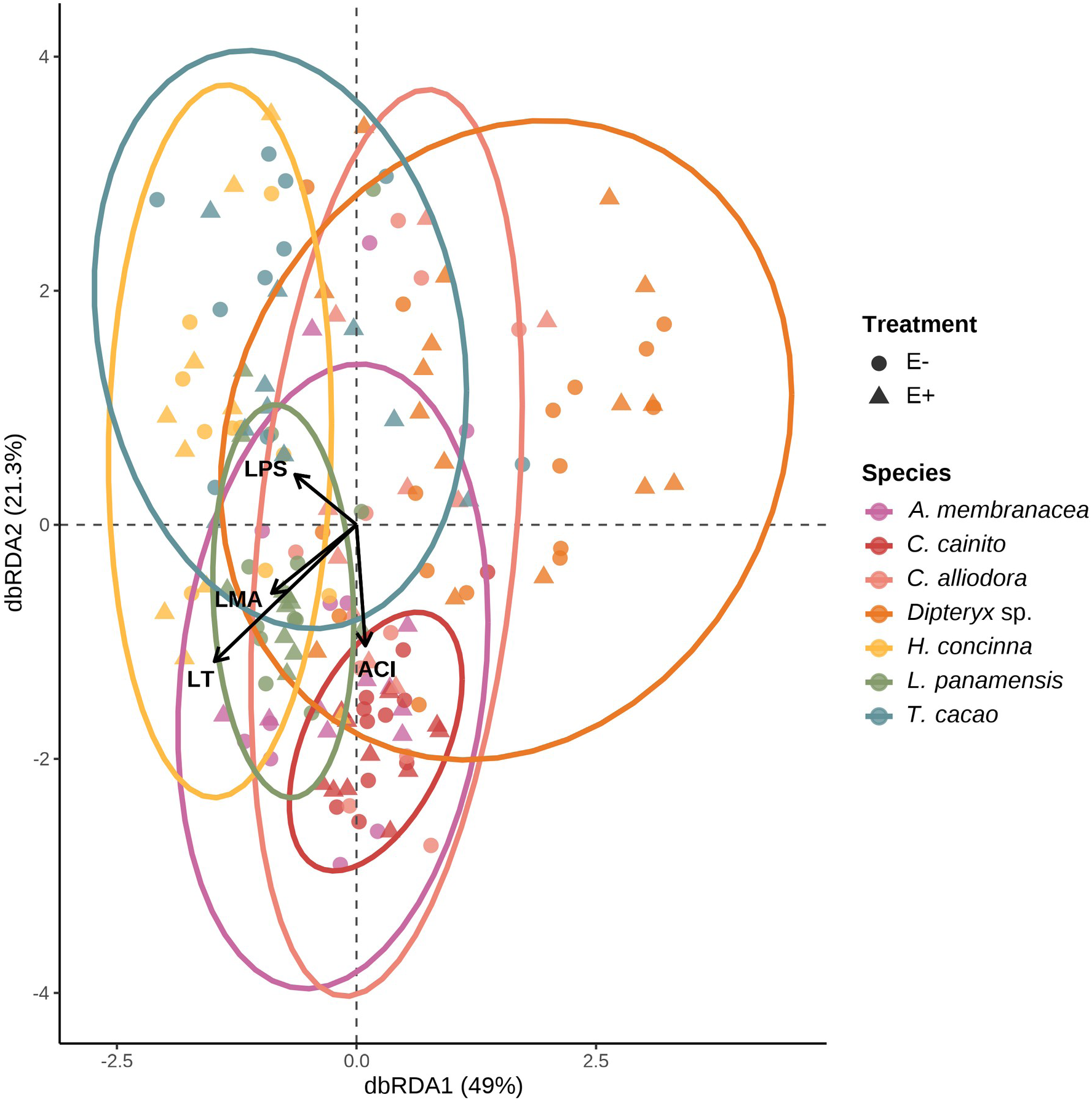
FEF community composition was associated with leaf functional traits from the leaf economic spectrum (genomic data set). variation in FEF community composition within and between host species (*n* = 7) and treatment groups (*E*-, *E*+) from distanmce-based redundancy analyses (dbRDA). Solid arrows represent statistically sisgnificant associations (*p* < .05). Each point represent a FEF community samples from one host tree species per treatment group; colors represent host tree species. Circles and filled triangles represent low (*E*-) and high (*E*+) FEF treatment groups, respectively.

There was a core set of OTUs that significantly associated with host tree species or particular treatments. In summary, 72 out of 569 Ascomycota OTUs were significantly associated with all host tree species (Table S5). Seedlings used in herbivory assays had 13 OTUs significantly associated with high herbivory damage (>70%) and 3 and 1 OTUs significantly associated with medium (31-69%) and low (<30%) herbivory damage, respectively, in *E-* and *E+* treatment groups (Table S6). We detected 11 OTUs significantly associated with seedlings that experienced high (>30%) pathogen damage in *E-* and *E+* treatment groups (Table S7). We could not tease apart which OTUs were associated with leaves exposed to *Calonectria* sp. agar plugs and non-exposed leaves because the genomic data scale for these trials was at the individual plant level, not leaf level. We found 30 OTUs significantly associated with *E+* treated (inoculated with forest spore fall) individuals used in our pathogen assays (Table S8).

Host species leaf traits encompassed variation along the leaf economic spectrum (LES). The PCA revealed how ACI, LT, LPS and LMA were related. We plotted leaf trait data according to tree species groups on the PCA axes to show how the variance in the complete data set is explained by PC1 (60%) and PC2 (27%) (Fig. 4a). We observed that ACI, LPS and LMA loadings tracked along PC1 towards more negative values, showing correlation among these traits (Fig. 4a). Traits LT and LPS were orthogonal to each other (Fig. 4a), indicative of low correlation. We note distinct grouping of host species along PC1 towards negative values. We saw compact clustering of host species on opposite ends of PC2 (Fig. 4a). We note similar relationships between the leaf traits with respect to PC1 and PC2 in the subset of individual seedlings used for herbivory versus pathogen damage trials (Fig. 4b-c). The PCA of leaf traits from seedlings used in herbivory trials had a PC1 explaining 57.5% of the variance and a PC2 explaining 28% of the variance in the subset data (Fig. 4b). We saw an inversion of the LT loading in direction of positive values, as well as the same tree species clustered (i.e., *Dypteryx* sp. and *A. membranacea*) along PC2 (Fig. 4b) with respect to Fig. 4a. The PCA of leaf traits from seedling used in pathogen damage trials had a PC1 explaining 64% of the variance and a PC2 explaining 25% of the variance in the subset data (Fig. 4c). We detected similar relationships among leaf traits and PC axes in the pathogen damage subset data (Fig. 4c) when compared to the complete data set (Fig. 4a).

**Figure 4.**
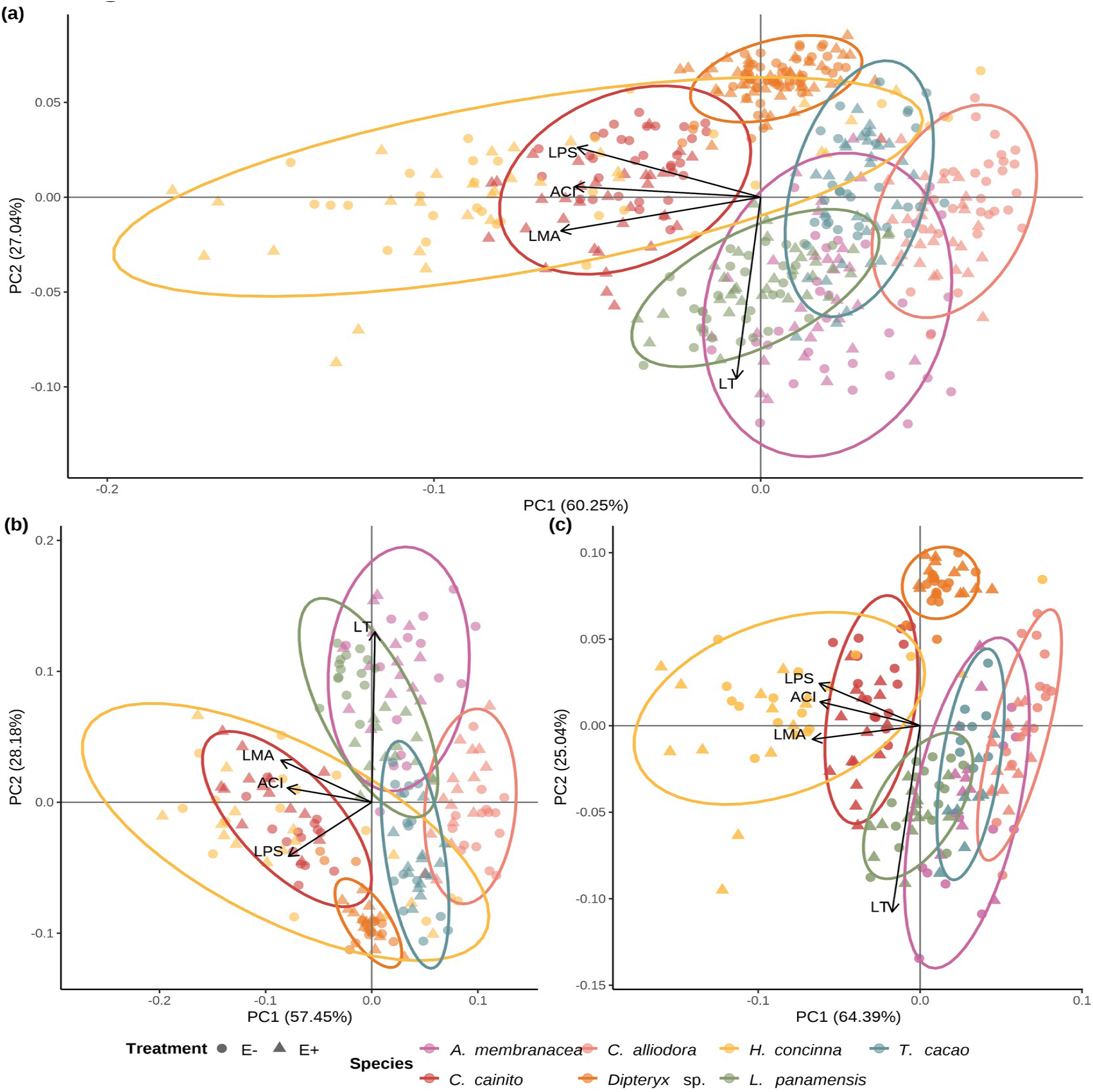
Leaf Functional traits are conserved within tree species regardless of endophyte load treatment. (a) Principal Component Analysis (PCA) of leaf functional traits from all tree species separated by *E-* and *E+* treatment. (b) PCA of leaf functional traits of plants solely used in ant herbivory assays. (c) PCA leaf functional traits of plants used solely in pathogen damage assays. Colors represent tree species. Circle and triangles represent low (*E-*) and high (*E+*) FEF treatment groups, respectively. Colored ellipses correspond to tree species and represent 95% confidence intervals.

Leaf functional traits influence plants’ response to herbivory and pathogen damage differentially. We examined the relationship between leaf functional traits, herbivory and pathogen damage to understand how host species on opposite ends of the LES modulate herbivory and pathogen damage. We used simple linear regressions plotting herbivory and pathogen damage with PC1 and PC2 from the PCA. Simple linear regressions of herbivory (%) against PC1 revealed no correlation (Fig. 5a; *R^2^_adj_*= -0.0024, *F*∼1, 208∼ = 0.508, *df* = 208, *p* = .447), where positive values represent greater values of ACI, LPS and LMA. Even though we note large spread in the data (Fig. 5a and 5b), we see that herbivory was strongly associated with PC2 (Fig. 5b; *R^2^_adj_* = 0.079, *F*∼1, 208∼ = 18.9, *p* < .0001), where positive values represent greater LT . Regressions of pathogen damage (%) plotted against PC1 revealed a significant correlation (Fig. 5c; *R^2^_adj_* = 0.064, *F*∼1, 380∼ = 26.93 *p* < .0001), in which positive values represent greater values of ACI, LPS and LMA. We did not see a significant relationship between pathogen damage and PC2 (Fig. 5d; *R^2^_adj_* = 0.002, *F*∼1, 380∼ = 1.60, *p* = .207). We uncovered similar patterns when we performed simple linear regressions on the raw leaf functional traits and logit transformed herbivory and pathogen damage data (Fig. S7 and S8, respectively). We observed a significant positive relationship between herbivory and LT (Figure S7a; *R^2^_adj_* = 0.081, *F*∼1, 208∼) = 19.45, *p* < .0001) when considering the complete data set . We did not observe a significant relationship between herbivory (%) and LPS, ACI, LMA, and Shannon diversity index for FEF (Figure S7b - S7e) for the complete data set. However, we did observe a general decline in herbivory as FEF diversity increased which aligns with our first prediction. Furthermore, we see a significant negative relationship between herbivory and Shannon diversity for the *E-* treatment group (Figure S7e; *R^2^_adj_* = 0.138, *F*∼1, 103∼ = 17.7, *p* < .0001). We also saw an increase in herbivory for the *E+* treatment group as Shannon diversity index for FEF increased, but this is not statistically significant (Figure S7e; *R^2^_adj_*= 0.024, *F*∼1, 103∼ = 3.55, *p* = 0.062). A result contrary to our expectations. We noted significant negative relationships between pathogen damage and LPS (Figure S8b; *R^2^_adj_* = 0.078, *F*∼1, 380∼ = 33.32, *p* < .0001), ACI (Figure S8c; *R^2^_adj_* = 0.033, *F*∼1, 380∼ = 14.34, *p* = .0002) and LMA (Figure S8d; *R^2^_adj_*= 0.030, *F*∼1, 380∼ = 12.6, *p* < .001) when considering the complete data set. Pathogen damage did not have a significant correlation with LT(Figure S8a; *R^2^_adj_* = -0.001, *F*∼1, 380∼ = 0.50, *p* = .482). The *E-* and *E+* treatment groups follow the same general trend as the complete data set. Contrary to our predictions, we found a statistically significant positive relationship between pathogen damage and Shannon diversity index in the complete data set (Figure S8e; *R^2^_adj_*= 0.015, *F*∼1, 380∼ = 6.90, *p* < .01). Upon further scrutiny, only the *E+* treatment group has a significant positive correlation between pathogen damage and Shannon diversity index (Figure S8e;*R^2^_adj_*= 0.031, *F*∼1, 188∼ = 7.11, *p* < .01).

**Figure 5.**
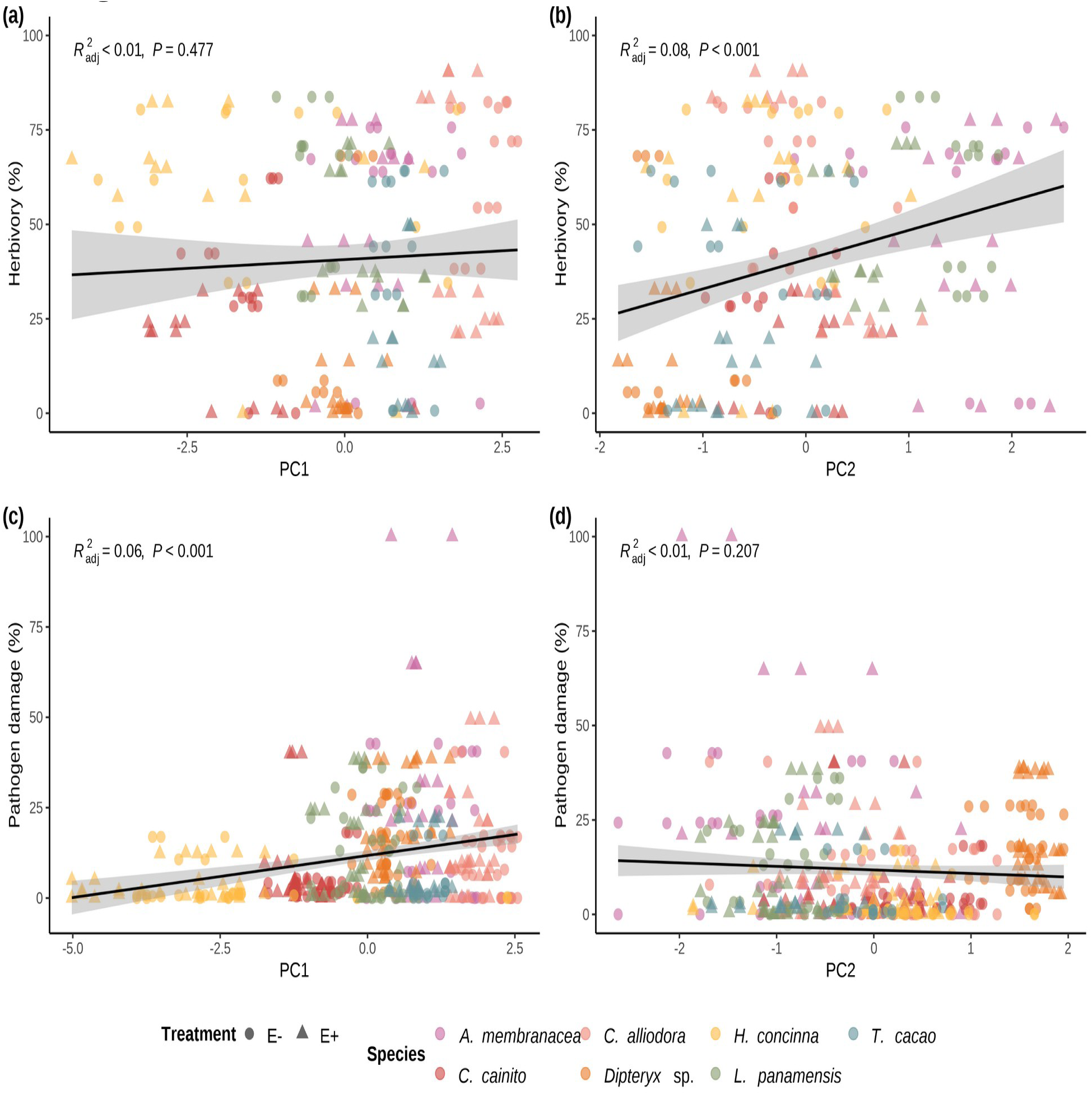
Simple linear regressions of herbivory and pathogen damage on PC1 and PC2 axes from PCAs of leaf traits for ant herbivory and pathogen damage assays. Linear regression of a) percent herbivory damage and PC1 axis (R^2^-adjusted= -0.0024, *p* = .447); b) percent herbivory damage and PC2 axis (R^2^-adjusted = 0.079, *p* = < .001); c) percent pathogen damage and PC1 axis (R^2^-adjusted = 0.064, *p* = < .001); and d) percent pathogen damage and PC2 axis (R^2^-adjusted = 0.0016, *p* = .207). Colors represent individual species. Circle and triangles represent *E-* and *E+* treatments, respectively.

Leaf thickness and LMA have opposite outcomes in plant’s response to herbivory. The best-fit for our GLMMs showed that fixed effects LT, LMA, and *E+* treatment group are statistically significant predictors of herbivory damage (Table 2). No measure of FEF abundance or diversity was present in the final model. Leaf mass per area is a significant positive predictor of herbivory with the greatest effect size (estimate = 1741, *t*(200) = 3.53 *p* < .001). While LT is a significant negative predictor of herbivory damage (estimate = -0.01, *t*(200) = -2.52, *p* = .01) and *E+* as well (estimate = -0.78, *t*(200) = -4.62, *p* < .001). The best fit model for pathogen damage did not reveal any of the leaf functional traits as significant predictors (Table 2). Like our previous model, no measure of FEF abundance or diversity was present in final model. Even though it was not significant, LMA showed the greatest effect size (estimate = 171.7, *t*(352) = 0.96, *p* = .34).

**Table 2:** Generalized linear mixed models (GLMMs) for predicting leaf herbivory and pathogen damage.

| Predictors | Herbivory model |  | Pathogen damage model |  |
| --- | --- | --- | --- | --- |
|  | Estimate | CI [t-statistic] | Estimate | CI [t-statistic] |
| *Fixed effects* |  |  |  |  |
| Intercept | -0.257 | (-2.315 - 1.801 ) [-0.246] | **-3.416 \*\*\*** | **(-4.403 - -2.428 ) [-6.801]** |
| Thickness | **-0.010 \*** | **(-0.018 - -0.002 ) [-2.517]** | 0.003 | (-0.001 - 0.007 ) [1.666] |
| LMA | **1741.216 \*\*\*** | **(768.816 - 2713.616 ) [3.531]** | 171.649 | (-181.400 - 524.698 ) [0.956] |
| Endophyte load (E+) | **-0.777 \*\*\*** | **(-1.109 - -0.445 ) [-4.619]** |  |  |
| *Random effects* |  |  |  |  |
| Intercept | 1.390 | (0.753 - 2.565 ) |  |  |
| Observations | 1.827 |  |  |  |
| *A. membranacea* | 1.000 |  | 1.000 |  |
| *C. cainito* | 0.654 |  | 0.780 |  |
| *C. alliodora* | 0.600 |  | 0.463 |  |
| *Dipteryx sp.* | 0.784 |  | 0.513 |  |
| *H. concinna* | 0.822 |  | 0.365 |  |
| *L. panamensis* | 0.425 |  | 0.312 |  |
| *T. cacao* | 0.782 |  | 0.767 |  |
| N | 210 |  | 382 |  |
| logLik | -358.893 |  | -492.620 |  |
| AIC | 741.785 |  | 1011.239 |  |

Significance levels are represented by asterisks [ $p < .05$ (\*), $p < .01$ (\*\*), $p < .001$ (\*\*\*), and $p < .0001$ (\*\*\*\*)]. T statistics in brackets.

## 6. Discussion

This is the first study to examine the interplay of FEF communities and leaf functional traits in response to the effects of generalist herbivore and pathogen in tropical tree species. Integrating the role FEF communities into a conceptual framework that includes trade-offs to plants’ constitutive and induced defenses in response to natural enemies help us understand the importance of FEF communities in the maintenance of plant diversity in tropical forests. Our findings suggest that foliar endophytic fungi (FEF) communities improve and leaf traits work to improve leaf defenses against generalist herbivores and pathogens in tropical trees. Across all host species, we saw that the *E+* treatment group exhibited lower herbivory damage than the *E-* treatment group. At the host species level, we saw that *E+* had less herbivory damage, however these differences were not significant between treatment groups. This is probably due to a small replicate size in treated individuals per species used in herbivory assay trials. Our results align with Estrada et al., (2013), where leaf-cutter ant effects were significantly reduced in paper disks and leaves treated with higher densities of a common endophyte. In a laboratory setting, Bittleston et al., (2011) found similar patterns in herbivory reduction using *C. alliodora* treated with high and low FEF loads and laboratory reared colonies of *Atta colombica*. In a field study, Coblentz and Van Bael (2013) found that leaf-cutter ants preferred leaves with a lower FEF abundance when compared to surrounding leaf material (material not selected by ants). Rocha et al. (2017), found endophytic *Trichoderma* more frequently isolated from leaf cuttings rejected by queen-less leaf-cutter ant colonies of *A. sexdens rubropilosa* in southeast Brazil. Our results track findings from previous studies that focus on single host-endophyte interactions or haphazard field collection and build upon them by using multiple host tree species and field inoculated FEF communities.

Our GLMM analysis allowed us to look at FEF and leaf traits at the same time; for herbivory damage we observed only LMA, LT and *E+* treatment group as strong predictors (Table 2). Using data from all the assays in simple linear regressions, FEF diversity and community composition correlated negatively with herbivory damage, since we saw reduced herbivory as Shannon diversity index increased (black regression line in Figure S7e). As a caveat to these results, we saw an inverse relationship between the treatment groups with the *E+* group positively correlating to herbivory and *E-* group negatively correlating to herbivory (yellow and pink regression lines, respectively, in Figure S7e). The contrast between GLMM and simple linear regressions point to the nuanced role of FEF diversity in combating herbivory; leaf functional traits, LMA and LT, overshadowed FEF diversity in our results. The FEF abundance, measured as *E+* treatment group, also played a strong role in predicting herbivory, but less so diversity and community composition.

We did not observe the same pattern for pathogen damage since there were no significant differences between treatment groups when considering all tree species combined (Figure 2b). Similar to herbivory damage, pathogen damage was not significantly different across treatment groups per species. Interestingly we saw that the *E+* group had equal or slightly higher pathogen damage than the *E-* group in all tree species (Figure S6b). The best-fit GLMM for pathogen damage showed LMA and LT as weak predictors (Table 2). On the other hand, with simple linear regressions we saw a significant increase in pathogen damage as Shannon diversity index increased for the complete data set (Figure S8e). Like our herbivory damage models, the contrast between GLMM and simple linear regression point to a nuanced role of FEF abundance and diversity in combating a pathogen. Pathogen damage significantly increased in the *E+* group as Shannon diversity increased, but the *E-* group did not (Figure S8e). González-Tauber (2016) found the opposite trend when examining *in vitro* the inhibitory effects of the most abundant genera (*Mycosphaerella* sp., *Xylaria* sp., *Diaporthe* sp., and *Penicillium* sp.) in FEF communities isolated from the southern temperate tree *Embothrium coccineum* against three common pathogens (*Botrytis cinerea*, *Fusarium oxysporum* and *Ceratocystis pilifera*). In our case, it is possible that the FEF abundance and diversity acquired from spore-fall have a synergistic interaction with *Calonectria* sp., thus outweighing any benefits provided by single-species interactions, ultimately increasing pathogen damage intensity.

The leaf economic spectrum (LES) allows us to interpret our results based on the functional traits that can have an important role in understanding tripartite plant-insect-pathogen interaction. The LES describes how leaf functional traits with high values are characteristic of long-lived versus short-lived leaves Poorter & Bongers (2006). The LES, thus acts as a host-imposed filter which FEF communities have to overcome to colonize leaf tissue, assemble, and carry out their life cycle. We did not measure leaf lifespan directly, but we interpret the values obtained for the leaf functional traits we measured as proxies for leaf lifespan, with the caveat that the traits were measured on seedlings and leaves were relatively young (< 150 days). We saw that species with relatively low values of ACI, LMA, and LPS (Figures S2 - S5) constrained in PC1 axis (Figure 4b) experienced very little herbivory damage (i.e., *Dypteryx* sp.) or had much variability (i.e., *C. alliodora*) (Figure 5a). Species with relatively high values of ACI, LMA, and LPS like *C. cainito* experienced low to moderate herbivory damage (Figure 5a). A similar pattern was evident in the pathogen damage data set (Figure 5c). The contrast in the slopes of the regression lines in Figure 5a and 5c highlight the potential importance LES traits have to counter herbivory and pathogen damage. We view these findings and those presented in Figure S7 as support for our prediction, where leaves on the high end of the LES, are less attractive to leaf cutter-ants but the effects of FEF communities outweigh this selection factor after a certain threshold is reached, as mentioned above.

Comprehending the relationship between the Leaf Economic Spectrum (LES), plant defenses, and FEF communities is crucial for understanding the complex interactions among plants, insects, and pathogens. The Optimal Defense Theory (ODT), as outlined by Stamp (2003), proposes three key predictions about plant defenses. First, a plant’s defense investment is directly proportional to the frequency of attacks, such as herbivory or pathogen intensity, and inversely related to the cost of resources (Holeski et al., 2010). Second, plants tend to allocate resources preferentially to parts with high reproductive value, especially when defense costs are minimal. Third, plants exhibit increased defensive responses after being attacked. This framework suggests that the likelihood of a plant to bolster its defenses following an attack is inversely related to its inherent defense traits (Holeski et al., 2010). Our results point to a preemptive low-cost investment strategy against plant enemies, particularly at the seedling stage, that is leveraged by species specific inherent defense mechanisms. We did not track herbivory or pathogen damage past the seedling stage, so an avenue for inquiry is to investigate how FEF communities change after herbivore and pathogen attacks and to determine the key life stages. A combination of *in vitro* and *in vivo* assays could help elucidate the roles of specific FEF OTUs. The use of *in vitro* assays could help identify the potential anti-herbivore and anti-pathogen qualities of specific FEF OTUs. *In vivo* inoculation assays could help identify the importance that specific FEF OTUs have in different developmental stages, especially in older stages.

## 7. Conclusion

This study advances our understanding of the intricate interactions between plants and their FEF communities, particularly in the context of plant defense mechanisms against herbivores and pathogens. Our findings highlight the complex dynamics of plant-herbivore-pathogen relationships and underscore the importance of FEF communities as a potentially low-cost, preemptive defense strategy for plants, especially during early growth stages. These insights not only shed light on the nuanced role of endophytes in plant ecology but also open avenues for future research, particularly in exploring the strategic resource allocation in plants and the specific contributions of FEF to plant resilience. As we continue to unravel these complex biological interactions, the knowledge gained holds promise not only for ecological theory but also for practical applications in agriculture, forestry, and conservation of the tropics.

## 8. Author Contributions

A.E.A., S.A.V. designed research study. M.S.J. and B.A.R. performed experiments and collected data. B.A.R. and M.S.J. analyzed data. B.A.R. wrote the manuscript with input from all authors. All authors gave final approval for publication. Our study included technicians based in the country where the study was carried out throughout the preparation phase of the project (seed collection and preparation). We recognize that more could have been done to engage local residents, students and scientists with our research as our project developed. We plan to address these caveats in future research

## 9. Acknowledgements

This research was funded by NSF DEB-1556583 to S.A.V. and NSF DEB-1556287 to A.E.A., and by the School of Plant Sciences and the College of Agriculture, Life and Environmental Sciences at The University of Arizona (AEA). We thank Ming-Min Lee and Nicole Colón-Carrion for laboratory support. We thank the Republic of Panama for the opportunity to conduct research there, and the Smithsonian Tropical Research Institute for logistics support.

## 10. Conflict of Interest Statement

The authors declare no competing interests.

## 11. Data Availability Statement

The genomic data that support the findings of this study will be submited to NCBI-GenBank upon acceptance of this manuscript. The manuscript and code for reproducibility is available from corresponding author’s GitHub.

## Notes

### Competing Interest Statement

The authors have declared no competing interest.

## References

1. Anderson, J. P., Gleason, C. A., Foley, R. C., Thrall, P. H., Burdon, J. B., & Singh, K. B. (2010). Plants versus pathogens: An evolutionary arms race. Functional Plant Biology, 37(6), 499. 10.1071/FP09304

2. Anderson, M. J. (2017). Permutational Multivariate Analysis of Variance (PERMANOVA). In Wiley StatsRef: Statistics Reference Online (pp. 1–15). Wiley. 10.1002/9781118445112.stat07841

3. Arnold, A. E., & Engelbrecht, B. M. J. (2007). Fungal endophytes nearly double minimum leaf conductance in seedlings of a neotropical tree species. Journal of Tropical Ecology, 23(3), 369–372. 10.1017/S0266467407004038

4. Arnold, A. E., Maynard, Z., Gilbert, G. S., Coley, P. D., & Kursar, T. A. (2000). Are tropical fungal endophytes hyperdiverse? Ecology Letters, 3(4), 267–274. 10.1046/j.1461-0248.2000.00159.x

5. Arnold, A. E., Mejía, L. C., Kyllo, D., Rojas, E. I., Maynard, Z., Robbins, N., & Herre, E. A. (2003). Fungal endophytes limit pathogen damage in a tropical tree. Proceedings of the National Academy of Sciences, 100(26), 15649–15654.

6. Bael, S. V., Estrada, C., & Arnold, A. E. (2017). Chapter 6 Foliar Endophyte Communities and Leaf Traits in Tropical Trees. In J. Dighton & J. F. White (Eds.), Mycology (pp. 79–94). CRC Press. 10.1201/9781315119496-7

7. Bartoń, K. (2023). *MuMIn: Multi-model inference* [Manual]. https://CRAN.R-project.org/package=MuMIn

8. Benjamini, Y., & Hochberg, Y. (1995). Controlling the False Discovery Rate: A Practical and Powerful Approach to Multiple Testing. Journal of the Royal Statistical Society: Series B (Methodological), 57(1), 289–300. 10.1111/j.2517-6161.1995.tb02031.x

9. Bittleston, L. S., Brockmann, F., Wcislo, W., & Van Bael, S. A. (2011). Endophytic fungi reduce leaf-cutting ant damage to seedlings. Biology Letters, 7(1), 30–32. 10.1098/rsbl.2010.0456

10. Blanchet, F. G., Legendre, P., & Borcard, D. (2008). FORWARD SELECTION OF EXPLANATORY VARIABLES. Ecology, 89(9), 2623–2632. 10.1890/07-0986.1

11. Callahan, B. J., McMurdie, P. J., Rosen, M. J., Han, A. W., Johnson, A. J. A., & Holmes, S. P. (2016). DADA2: High-resolution sample inference from Illumina amplicon data. Nature Methods, 13(7), 581–583. 10.1038/nmeth.3869

12. Carbone, I., White, J. B., Miadlikowska, J., Arnold, A. E., Miller, M. A., Kauff, F., U’Ren, J. M., May, G., & Lutzoni, F. (2017). T-BAS: Tree-Based Alignment Selector toolkit for phylogenetic-based placement, alignment downloads and metadata visualization: An example with the Pezizomycotina tree of life. Bioinformatics, 33(8), 1160–1168. 10.1093/bioinformatics/btw808

13. Carbone, I., White, J. B., Miadlikowska, J., Arnold, A. E., Miller, M. A., Magain, N., U’Ren, J. M., & Lutzoni, F. (2019). T-BAS Version 2.1: Tree-Based Alignment Selector Toolkit for Evolutionary Placement of DNA Sequences and Viewing Alignments and Specimen Metadata on Curated and Custom Trees. Microbiology Resource Announcements, 8(29), e00328–19. 10.1128/MRA.00328-19

14. Chagas, F. O., Pessotti, R. D. C., Caraballo-Rodríguez, A. M., & Pupo, M. T. (2018). Chemical signaling involved in plant–microbe interactions. Chemical Society Reviews, 47(5), 1652–1704. 10.1039/C7CS00343A

15. Christian, N., Whitaker, B. K., & Clay, K. (2017). Chapter 5 A Novel Framework for Decoding Fungal Endophyte Diversity. In J. Dighton & J. F. White (Eds.), Mycology (pp. 63–78). CRC Press. 10.1201/9781315119496-6

16. Coblentz, K. E., & Van Bael, S. A. (2013). Field colonies of leaf-cutting ants select plant materials containing low abundances of endophytic fungi. Ecosphere, 4(5). 10.1890/ES13-00012.1

17. Currie, A. F., Wearn, J., Hodgson, S., Wendt, H., Broughton, S., & Jin, L. (2014). Foliar Fungal Endophytes in Herbaceous Plants: A Marriage of Convinience? In V. C. Verma & A. C. Gange (Eds.), Advances in Endophytic Research (pp. 61–81). Springer India. 10.1007/978-81-322-1575-2

18. Daru, B. H., Bowman, E. A., Pfister, D. H., & Arnold, A. E. (2019). A novel proof of concept for capturing the diversity of endophytic fungi preserved in herbarium specimens. Philosophical Transactions of the Royal Society B: Biological Sciences, 374(1763), 20170395. 10.1098/rstb.2017.0395

19. De Cáceres, M., & Legendre, P. (2009). Associations between species and groups of sites: Indices and statistical inference. Ecology, 90, 3566–3574. 10.1890/08-1823.1

20. Estrada, C., Wcislo, W. T., & Van Bael, S. A. (2013). Symbiotic fungi alter plant chemistry that discourages leaf-cutting ants. New Phytologist, 198(1), 241–251. 10.1111/nph.12140

21. Feild, T. S., & Arens, N. C. (2005). Form, function and environments of the early angiosperms: Merging extant phylogeny and ecophysiology with fossils. New Phytologist, 166(2), 383–408. 10.1111/j.1469-8137.2005.01333.x

22. Fox, J., & Weisberg, S. (2019). An R companion to applied regression (3rd ed.). Sage. https://socialsciences.mcmaster.ca/jfox/Books/Companion/

23. Friesen, M. L., Porter, S. S., Stark, S. C., Von Wettberg, E. J., Sachs, J. L., & Martinez-Romero, E. (2011). Microbially Mediated Plant Functional Traits. Annual Review of Ecology, Evolution, and Systematics, 42(1), 23–46. 10.1146/annurev-ecolsys-102710-145039

24. Gilbert, G. S., & Webb, C. O. (2007). Phylogenetic signal in plant pathogen–host range. Proceedings of the National Academy of Sciences, 104(12), 4979–4983. 10.1073/pnas.0607968104

25. González-Teuber, M. (2016). The defensive role of foliar endophytic fungi for a South American tree. AoB PLANTS, 8, plw050. 10.1093/aobpla/plw050

26. Guerriero, G., Berni, R., Muñoz-Sanchez, J., Apone, F., Abdel-Salam, E., Qahtan, A., Alatar, A., Cantini, C., Cai, G., Hausman, J.-F., Siddiqui, K., Hernández-Sotomayor, S., & Faisal, M. (2018). Production of Plant Secondary Metabolites: Examples, Tips and Suggestions for Biotechnologists. Genes, 9(6), 309. 10.3390/genes9060309

27. Hanley, M. E., Lamont, B. B., Fairbanks, M. M., & Rafferty, C. M. (2007). Plant structural traits and their role in anti-herbivore defence. Perspectives in Plant Ecology, Evolution and Systematics, 8(4), 157–178. 10.1016/j.ppees.2007.01.001

28. Higgins, K. L., Arnold, A. E., Coley, P. D., & Kursar, T. A. (2014). Communities of fungal endophytes in tropical forest grasses: Highly diverse host- and habitat generalists characterized by strong spatial structure. Fungal Ecology, 8(1), 1–11. 10.1016/j.funeco.2013.12.005

29. Holeski, L. M., Chase-Alone, R., & Kelly, J. K. (2010). The genetics of phenotypic plasticity in plant defense: Trichome production in Mimulus guttatus. American Naturalist, 175(4), 391–400. 10.1086/651300

30. Kassambara, A. (2023a). *Ggpubr: ’ggplot2’ Based Publication Ready Plots* (R package version 0.6.0) [Computer software]. https://rpkgs.datanovia.com/ggpubr/

31. Kassambara, A. (2023b). *Rstatix: Pipe-Friendly Framework for Basic Statistical Tests* (R package version 0.7.2) [Computer software]. https://rpkgs.datanovia.com/rstatix/

32. Kitajima, K., Cordero, R. A., & Wright, S. J. (2013). Leaf life span spectrum of tropical woody seedlings: Effects of light and ontogeny and consequences for survival. Annals of Botany, 112(4), 685–699. 10.1093/aob/mct036

33. Kitajima, K., Llorens, A., Stefanescu, C., Timchenko, M. V., Lucas, P. W., & Wright, S. J. (2012). How cellulose-based leaf toughness and lamina density contribute to long leaf lifespans of shade-tolerant species. New Phytologist, 195(3), 640–652. 10.1111/j.1469-8137.2012.04203.x

34. Leakey, A. D. B., & Lau, J. A. (2012). Evolutionary context for understanding and manipulating plant responses to past, present and future atmospheric [CO _2_]. Philosophical Transactions of the Royal Society B: Biological Sciences, 367(1588), 613–629. 10.1098/rstb.2011.0248

35. Legendre, P., & Anderson, M. J. (1999). Distance-based redundancy analysis: Testing multispecies responses in multifactorial ecological experiments. Ecological Monographs, 69(1), 1–24. 10.1890/0012-9615(1999)069[0001:DBRATM]2.0.CO;2

36. Legendre, P., & Legendre, L. (2012). Numerical ecology (3d English edition). Elsevier.

37. Legendre, P., Oksanen, J., & Ter Braak, C. J. F. (2011). Testing the significance of canonical axes in redundancy analysis. Methods in Ecology and Evolution, 2(3), 269–277. 10.1111/j.2041-210X.2010.00078.x

38. Leigh, E. G., Rand, A. S., Windsor, D. M., & Institute, S. T. R. (Eds.). (1996). The ecology of a tropical forest: Seasonal rhythms and long-term changes (2nd ed). Smithsonian Institution Press.

39. Mason, C. M., & Donovan, L. A. (2015). Does investment in leaf defenses drive changes in leaf economic strategy? A focus on whole-plant ontogeny. Oecologia, 177(4), 1053–1066. 10.1007/s00442-014-3177-2

40. McArdle, B. H., & Anderson, M. J. (2001). FITTING MULTIVARIATE MODELS TO COMMUNITY DATA: A COMMENT ON DISTANCE-BASED REDUNDANCY ANALYSIS. Ecology, 82(1), 290–297. 10.1890/0012-9658(2001)082[0290:FMMTCD]2.0.CO;2

41. McGill, B. J., Enquist, B. J., Weiher, E., & Westoby, M. (2006). Rebuilding community ecology from functional traits. Trends in Ecology and Evolution. 10.1016/j.tree.2006.02.002

42. McMurdie, P. J., & Holmes, S. (2013). Phyloseq: An R Package for Reproducible Interactive Analysis and Graphics of Microbiome Census Data. PLoS ONE, 8(4), e61217. 10.1371/journal.pone.0061217

43. Mejía, L. C., Herre, E. A., Sparks, J. P., Winter, K., García, M. N., Van Bael, S. A., Stitt, J., Shi, Z., Zhang, Y., Guiltinan, M. J., & Maximova, S. N. (2014). Pervasive effects of a dominant foliar endophytic fungus on host genetic and phenotypic expression in a tropical tree. Frontiers in Microbiology, 5, 1–16. 10.3389/fmicb.2014.00479

44. Mejía, L. C., Rojas, E. I., Maynard, Z., Bael, S. V., Arnold, A. E., Hebbar, P., Samuels, G. J., Robbins, N., & Herre, E. A. (2008). Endophytic fungi as biocontrol agents of Theobroma cacao pathogens. Biological Control, 46(1), 4–14. 10.1016/j.biocontrol.2008.01.012

45. Niklas, K. J., Shi, P., Gielis, J., Schrader, J., & Niinemets, Ü. (2023). Editorial: Leaf functional traits: Ecological and evolutionary implications. Frontiers in Plant Science, 14, 1169558. 10.3389/fpls.2023.1169558

46. Oita, S., Ibáñez, A., Lutzoni, F., Miadlikowska, J., Geml, J., Lewis, L. A., Hom, E. F. Y., Carbone, I., U’Ren, J. M., & Arnold, A. E. (2021). Climate and seasonality drive the richness and composition of tropical fungal endophytes at a landscape scale. Communications Biology, 4(1), 313. 10.1038/s42003-021-01826-7

47. Oksanen, J., Simpson, G. L., Blanchet, F. G., Kindt, R., Legendre, P., Minchin, P. R., O’Hara, R. B., Solymos, P., Stevens, M. H. H., Szoecs, E., Wagner, H., Barbour, M., Bedward, M., Bolker, B., Borcard, D., Carvalho, G., Chirico, M., De Caceres, M., Durand, S., … Weedon, J. (2022). Vegan: Community Ecology Package. https://github.com/vegandevs/vegan

48. Pinheiro, J. C., & Bates, D. M. (2000). Mixed-Effects Models in S and S-PLUS. Springer. 10.1007/b98882

49. Pinheiro, J., Bates, D., & R Core Team. (2023). Nlme: Linear and nonlinear mixed effects models [Manual]. https://CRAN.R-project.org/package=nlme

50. Poorter, L., & Bongers, F. (2006). LEAF TRAITS ARE GOOD PREDICTORS OF PLANT PERFORMANCE ACROSS 53 RAIN FOREST SPECIES. Ecology, 87(7), 1733–1743. 10.1890/0012-9658(2006)87[1733:LTAGPO]2.0.CO;2

51. Porras-Alfaro, A., & Bayman, P. (2011). Hidden Fungi, Emergent Properties: Endophytes and Microbiomes. Annual Review of Phytopathology, 49(1), 291–315. 10.1146/annurev-phyto-080508-081831

52. R Core Team. (2024). R: A language and environment for statistical computing [Manual]. R Foundation for Statistical Computing. https://www.R-project.org/

53. Rocha, S. L., Evans, H. C., Jorge, V. L., Cardoso, L. A. O., Pereira, F. S. T., Rocha, F. B., Barreto, R. W., Hart, A. G., & Elliot, S. L. (2017). Recognition of endophytic *Trichoderma* species by leaf-cutting ants and their potential in a Trojan-horse management strategy. Royal Society Open Science, 4(4), 160628. 10.1098/rsos.160628

54. Rodriguez, R. J., White, J. F., Arnold, A. E., & Redman, R. S. (2009). Fungal endophytes: Diversity and functional roles. New Phytologist, 182(2), 314–330. 10.1111/j.1469-8137.2009.02773.x

55. Rognes, T., Flouri, T., Nichols, B., Quince, C., & Mahé, F. (2016). VSEARCH: A versatile open source tool for metagenomics. PeerJ, 4, e2584. 10.7717/peerj.2584

56. Sarmiento, C., Zalamea, P. C., Dalling, J. W., Davis, A. S., Simon, S. M., U’Ren, J. M., & Arnold, A. E. (2017). Soilborne fungi have host affinity and host-specific effects on seed germination and survival in a lowland tropical forest. Proceedings of the National Academy of Sciences of the United States of America, 114(43), 11458–11463. 10.1073/pnas.1706324114

57. Saunders, M., Glenn, A. E., & Kohn, L. M. (2010). Exploring the evolutionary ecology of fungal endophytes in agricultural systems: Using functional traits to reveal mechanisms in community processes. Evolutionary Applications, 3(5-6), 525–537. 10.1111/j.1752-4571.2010.00141.x

58. Schneider, C. A., Rasband, W. S., & Eliceiri, K. W. (2012). NIH Image to ImageJ: 25 years of image analysis. Nature Methods, 9(7), 671–675. 10.1038/nmeth.2089

59. Stamp, N. (2003). Out of the quagmire of plant defense hypotheses. Quarterly Review of Biology, 78(1), 23–55. 10.1086/367580

60. Tellez, P. H., Arnold, A. E., Leo, A. B., Kitajima, K., & Van Bael, S. A. (2022). Traits along the leaf economics spectrum are associated with communities of foliar endophytic symbionts. Frontiers in Microbiology, 13, 927780. https://doi.org/lutzoni

61. Tellez, P. H., Rojas, E., & Van Bael, S. (2016). Red coloration in young tropical leaves associated with reduced fungal pathogen damage. Biotropica, 48(2), 150–153. 10.1111/btp.12303

62. Teoh, E. S. (2016). Secondary Metabolites of Plants. In E. S. Teoh, Medicinal Orchids of Asia (pp. 59–73). Springer International Publishing. 10.1007/978-3-319-24274-3_5

63. U’Ren, J. M., & Arnold, A. E. (2017). 96 well DNA Extraction Protocol for Plant and Lichen Tissue Stored in CTAB. Protocols.io, 1–5. 10.17504/protocols.io.fscbnaw

64. U’Ren, J. M., Lutzoni, F., Miadlikowska, J., Zimmerman, N. B., Carbone, I., May, G., & Arnold, A. E. (2019). Host availability drives distributions of fungal endophytes in the imperilled boreal realm. Nature Ecology & Evolution, 3(10), 1430–1437. 10.1038/s41559-019-0975-2

65. Weiss, S., Xu, Z. Z., Peddada, S., Amir, A., Bittinger, K., Gonzalez, A., Lozupone, C., Zaneveld, J. R., Vázquez-Baeza, Y., Birmingham, A., Hyde, E. R., & Knight, R. (2017). Normalization and microbial differential abundance strategies depend upon data characteristics. Microbiome, 5(1), 27. 10.1186/s40168-017-0237-y

66. Wright, I. J., Reich, P. B., Cornelissen, J. H. C., Falster, D. S., Garnier, E., Hikosaka, K., Lamont, B. B., Lee, W., Oleksyn, J., Osada, N., Poorter, H., Villar, R., Warton, D. I., & Westoby, M. (2005). Assessing the generality of global leaf trait relationships. New Phytologist, 166(2), 485–496. 10.1111/j.1469-8137.2005.01349.x

67. Wright, I. J., Reich, P. B., Westoby, M., Ackerly, D. D., Baruch, Z., Bongers, F., Cavender-Bares, J., Chapin, T., Cornelissen, J. H. C., Diemer, M., Flexas, J., Garnier, E., Groom, P. K., Gulias, J., Hikosaka, K., Lamont, B. B., Lee, T., Lee, W., Lusk, C., … Villar, R. (2004). The worldwide leaf economics spectrum. Nature, 428(6985), 821–827. 10.1038/nature02403

