## Supplemental_Materials for "Evaluating the Role of Endophyte-Rich Leaves in Protecting Tropical Trees Against a Generalist Herbivore and a Pathogen"

### Supplementary Materials

#### Table S1: Student's t-Tests of mean anthocyanins (ACI)

Pairwise comparisons of ACI between species.

| | *p* - values | | | |
|---|---|---|---|---|
| **Comparison Species[1]** | **p** | **p.adj** | **p.format** | **p.signif[2]** |
| ***A. membranacea*** | | | | |
| *C. cainito* | **2.636 × 10^-16** | **4.500 × 10^-15** | **0** | **** |
| *C. alliodora* | **3.713 × 10^-15** | **5.600 × 10^-14** | **0** | **** |
| *Dipteryx* sp. | **2.296 × 10^-6** | **1.600 × 10^-5** | **0** | **** |
| *H. concinna* | **6.179 × 10^-6** | **3.700 × 10^-5** | **0** | **** |
| *L. panamensis* | **1.538 × 10^-2** | **4.600 × 10^-2** | **0.02** | * |
| *T. cacao* | **9.137 × 10^-8** | **8.200 × 10^-7** | **0** | **** |
| ***C. cainito*** | | | | |
| *C. alliodora* | **3.154 × 10^-23** | **6.300 × 10^-22** | **< 2e-16** | **** |
| *Dipteryx* sp. | **4.559 × 10^-10** | **5.000 × 10^-9** | **0** | **** |
| *H. concinna* | 5.309 × 10^-1 | 5.300 × 10^-1 | 0.53 | ns |
| *L. panamensis* | **1.599 × 10^-11** | **2.200 × 10^-10** | **0** | **** |
| *T. cacao* | **3.656 × 10^-22** | **6.900 × 10^-21** | **< 2e-16** | **** |
| ***C. alliodora*** | | | | |
| *Dipteryx* sp. | **1.150 × 10^-26** | **2.400 × 10^-25** | **< 2e-16** | **** |
| *H. concinna* | **7.276 × 10^-11** | **9.500 × 10^-10** | **0** | **** |
| *L. panamensis* | **2.486 × 10^-15** | **4.000 × 10^-14** | **0** | **** |
| *T. cacao* | **1.428 × 10^-7** | **1.100 × 10^-6** | **0** | **** |
| ***Dipteryx* sp.** | | | | |
| *H. concinna* | **3.050 × 10^-3** | **1.200 × 10^-2** | **0** | ** |
| *L. panamensis* | 6.646 × 10^-2 | 1.300 × 10^-1 | 0.07 | ns |
| *T. cacao* | **3.807 × 10^-18** | **6.900 × 10^-17** | **< 2e-16** | **** |
| ***H. concinna*** | | | | |
| *L. panamensis* | **3.062 × 10^-4** | **1.500 × 10^-3** | **0** | *** |
| *T. cacao* | **5.704 × 10^-9** | **5.700 × 10^-8** | **0** | **** |
| ***L. panamensis*** | | | | |
| *T. cacao* | **3.583 × 10^-10** | **4.300 × 10^-9** | **0** | **** |

[1] *n* = 156 individuals

[2]Significance levels are represented by *ast*erisks [*p* < .05 (*), *p* <= .01 (**), *p* <= .001 (***), and *p* < .0001 (****)].

#### Table S2: Student's t-Tests of mean leaf thickness (LT) (µm)

Pairwise comparisons of LT between species.

| | *p* - values | | | |
|---|---|---|---|---|
| **Comparison Species**[1] | **p** | **p.adj** | **p.format** | **p.signif**[2] |
| ***A. membranacea*** | | | | |
| *C. cainito* | **1.793 × 10^-8** | **2.300 × 10^-7** | **0** | **** |
| *C. alliodora* | **6.857 × 10^-8** | **8.200 × 10^-7** | **0** | **** |
| *Dipteryx* sp. | **7.836 × 10^-18** | **1.500 × 10^-16** | **< 2e-16** | **** |
| *H. concinna* | **1.255 × 10^-5** | **1.000 × 10^-4** | **0** | **** |
| *L. panamensis* | 4.986 × 10^-1 | 1 | 0.5 | ns |
| *T. cacao* | **1.604 × 10^-6** | **1.400 × 10^-5** | **0** | **** |
| ***C. cainito*** | | | | |
| *C. alliodora* | 7.854 × 10^-1 | 1 | 0.79 | ns |
| *Dipteryx* sp. | **2.605 × 10^-19** | **5.200 × 10^-18** | **< 2e-16** | **** |
| *H. concinna* | 6.876 × 10^-2 | 4.800 × 10^-1 | 0.07 | ns |
| *L. panamensis* | **1.382 × 10^-12** | **2.100 × 10^-11** | **0** | **** |
| *T. cacao* | 4.765 × 10^-1 | 1 | 0.48 | ns |
| ***C. alliodora*** | | | | |
| *Dipteryx* sp. | **8.662 × 10^-17** | **1.500 × 10^-15** | **< 2e-16** | **** |
| *H. concinna* | 1.347 × 10^-1 | 8.100 × 10^-1 | 0.14 | ns |
| *L. panamensis* | **1.161 × 10^-10** | **1.600 × 10^-9** | **0** | **** |
| *T. cacao* | 6.481 × 10^-1 | 1 | 0.65 | ns |
| ***Dipteryx* sp.** | | | | |
| *H. concinna* | **8.177 × 10^-17** | **1.500 × 10^-15** | **< 2e-16** | **** |
| *L. panamensis* | **8.008 × 10^-32** | **1.700 × 10^-30** | **< 2e-16** | **** |
| *T. cacao* | **4.639 × 10^-13** | **7.400 × 10^-12** | **0** | **** |
| ***H. concinna*** | | | | |
| *L. panamensis* | **7.274 × 10^-7** | **7.300 × 10^-6** | **0** | **** |
| *T. cacao* | 3.649 × 10^-1 | 1 | 0.37 | ns |
| ***L. panamensis*** | | | | |
| *T. cacao* | **1.707 × 10^-7** | **1.900 × 10^-6** | **0** | **** |

[1] *n* = 156 individuals

[2]Significance levels are represented by *ast*erisks [*p* < .05 (*), *p* <= .01 (**), *p* <= .001 (***), and *p* < .0001 (****)].

#### Table S3: Student's t-Tests of mean leaf punch strength (LPS) (N mm$^{-1}$)

Pairwise comparisons of LPS between species.

| | *p* - values | | | |
|---|---|---|---|---|
| **Comparison Species[1]** | **p** | **p.adj** | **p.format** | **p.signif[2]** |
| ***A. membranacea*** | | | | |
| *C. cainito* | **9.032 × 10^-36** | **1.600 × 10^-34** | **< 2e-16** | **** |
| *C. alliodora* | 3.180 × 10^-1 | 3.200 × 10^-1 | 0.32 | ns |
| *Dipteryx* sp. | **3.538 × 10^-43** | **7.400 × 10^-42** | **< 2e-16** | **** |
| *H. concinna* | **7.548 × 10^-21** | **8.700 × 10^-20** | **< 2e-16** | **** |
| *L. panamensis* | **7.304 × 10^-26** | **1.200 × 10^-24** | **< 2e-16** | **** |
| *T. cacao* | **7.242 × 10^-21** | **8.700 × 10^-20** | **< 2e-16** | **** |
| ***C. cainito*** | | | | |
| *C. alliodora* | **3.873 × 10^-39** | **7.700 × 10^-38** | **< 2e-16** | **** |
| *Dipteryx* sp. | **3.649 × 10^-16** | **2.200 × 10^-15** | **0** | **** |
| *H. concinna* | **6.101 × 10^-12** | **2.400 × 10^-11** | **0** | **** |
| *L. panamensis* | **3.975 × 10^-28** | **6.800 × 10^-27** | **< 2e-16** | **** |
| *T. cacao* | **7.651 × 10^-21** | **8.700 × 10^-20** | **< 2e-16** | **** |
| ***C. alliodora*** | | | | |
| *Dipteryx* sp. | **1.738 × 10^-36** | **3.300 × 10^-35** | **< 2e-16** | **** |
| *H. concinna* | **1.267 × 10^-21** | **1.600 × 10^-20** | **< 2e-16** | **** |
| *L. panamensis* | **8.205 × 10^-21** | **8.700 × 10^-20** | **< 2e-16** | **** |
| *T. cacao* | **1.371 × 10^-22** | **1.900 × 10^-21** | **< 2e-16** | **** |
| ***Dipteryx* sp.** | | | | |
| *H. concinna* | **1.617 × 10^-15** | **8.100 × 10^-15** | **0** | **** |
| *L. panamensis* | **7.768 × 10^-26** | **1.200 × 10^-24** | **< 2e-16** | **** |
| *T. cacao* | **3.965 × 10^-7** | **8.800 × 10^-7** | **0** | **** |
| ***H. concinna*** | | | | |
| *L. panamensis* | **2.293 × 10^-18** | **1.800 × 10^-17** | **< 2e-16** | **** |
| *T. cacao* | **1.173 × 10^-17** | **8.200 × 10^-17** | **< 2e-16** | **** |
| ***L. panamensis*** | | | | |
| *T. cacao* | **2.949 × 10^-7** | **8.800 × 10^-7** | **0** | **** |

[1] *n* = 156 individuals

[2]Significance levels are represented by *ast*erisks [*p* < .05 (*), *p* <= .01 (**), *p* <= .001 (***), and *p* < .0001 (****)].

#### Table S4: Student's t-Tests of mean leaf mass per area (LMA) (mg $mm^{-2}$)

Pairwise comparisons of LMA between species.

| | *p* - values | | | |
|---|---|---|---|---|
| **Comparison Species[1]** | **p** | **p.adj** | **p.format** | **p.signif[2]** |
| ***A. membranacea*** | | | | |
| *C. cainito* | **9.032 × 10^-36** | **1.600 × 10^-34** | **< 2e-16** | **** |
| *C. alliodora* | 3.180 × 10^-1 | 3.200 × 10^-1 | 0.32 | ns |
| *Dipteryx* sp. | **3.538 × 10^-43** | **7.400 × 10^-42** | **< 2e-16** | **** |
| *H. concinna* | **7.548 × 10^-21** | **8.700 × 10^-20** | **< 2e-16** | **** |
| *L. panamensis* | **7.304 × 10^-26** | **1.200 × 10^-24** | **< 2e-16** | **** |
| *T. cacao* | **7.242 × 10^-21** | **8.700 × 10^-20** | **< 2e-16** | **** |
| ***C. cainito*** | | | | |
| *C. alliodora* | **3.873 × 10^-39** | **7.700 × 10^-38** | **< 2e-16** | **** |
| *Dipteryx* sp. | **3.649 × 10^-16** | **2.200 × 10^-15** | **0** | **** |
| *H. concinna* | **6.101 × 10^-12** | **2.400 × 10^-11** | **0** | **** |
| *L. panamensis* | **3.975 × 10^-28** | **6.800 × 10^-27** | **< 2e-16** | **** |
| *T. cacao* | **7.651 × 10^-21** | **8.700 × 10^-20** | **< 2e-16** | **** |
| ***C. alliodora*** | | | | |
| *Dipteryx* sp. | **1.738 × 10^-36** | **3.300 × 10^-35** | **< 2e-16** | **** |
| *H. concinna* | **1.267 × 10^-21** | **1.600 × 10^-20** | **< 2e-16** | **** |
| *L. panamensis* | **8.205 × 10^-21** | **8.700 × 10^-20** | **< 2e-16** | **** |
| *T. cacao* | **1.371 × 10^-22** | **1.900 × 10^-21** | **< 2e-16** | **** |
| ***Dipteryx* sp.** | | | | |
| *H. concinna* | **1.617 × 10^-15** | **8.100 × 10^-15** | **0** | **** |
| *L. panamensis* | **7.768 × 10^-26** | **1.200 × 10^-24** | **< 2e-16** | **** |
| *T. cacao* | **3.965 × 10^-7** | **8.800 × 10^-7** | **0** | **** |
| ***H. concinna*** | | | | |
| *L. panamensis* | **2.293 × 10^-18** | **1.800 × 10^-17** | **< 2e-16** | **** |
| *T. cacao* | **1.173 × 10^-17** | **8.200 × 10^-17** | **< 2e-16** | **** |
| ***L. panamensis*** | | | | |
| *T. cacao* | **2.949 × 10^-7** | **8.800 × 10^-7** | **0** | **** |

[1] *n* = 156 individuals

[2]Significance levels are represented by *ast*erisks [$p < .05$ (*), $p <= .01$ (**), $p <= .001$ (***), and $p < .0001$ (****)].

#### Table S5: Taxonomy of OTUs significantly correlated OTUs with tree host species.

| | | | | | | | | Multilevel pattern analysis | | | |
|---|---|---|---|---|---|---|---|---|---|---|---|
| **Kingdom** | **Phylum** | **Class** | **Order** | **Family** | **Genus** | **Species** | **OTU** | **Index** | **Stat** | ***p*[1]** | ***p*adj[2]** |
| ***T. cacao*** | | | | | | | | | | | |
| Fungi | Ascomycota | Dothideomycetes | Capnodiales | Mycosphaerellaceae | Coremiopassalora | *Coremiopassalora leptophlebae* | OTU 2 | 7 | 0.51 | ** | 0.01 |
| Fungi | Ascomycota | Dothideomycetes | Capnodiales | Mycosphaerellaceae | Pseudocercospora | *Pseudocercospora sp* | OTU 5 | 7 | 0.53 | ** | 0.01 |
| Fungi | Ascomycota | Eurotiomycetes | Eurotiales | Trichocomaceae | Talaromyces | *Talaromyces sp* | OTU 3 | 7 | 0.47 | ** | 0.01 |
| Fungi | Ascomycota | unidentified | unidentified | unidentified | unidentified | *Ascomycota sp* | OTU 14 | 7 | 0.5 | ** | 0.01 |
| Fungi | Ascomycota | Dothideomycetes | Capnodiales | Cladosporiaceae | Cladosporium | *Cladosporium sp* | OTU 20 | 7 | 0.38 | ** | 0.01 |
| Fungi | Ascomycota | Dothideomycetes | Capnodiales | Mycosphaerellaceae | Zasmidium | *Zasmidium queenslandicum* | OTU 95 | 7 | 0.39 | ** | 0.01 |
| Fungi | Ascomycota | Sordariomycetes | Hypocreales | Hypocreales_fam_Incertae_sedis | Acremonium | *Acremonium sp* | OTU 84 | 7 | 0.43 | ** | 0.01 |
| Fungi | Ascomycota | Dothideomycetes | Capnodiales | Dissoconiaceae | Ramichloridium | *Ramichloridium sp* | OTU 79 | 7 | 0.36 | ** | 0.02 |
| Fungi | Ascomycota | Dothideomycetes | Capnodiales | Mycosphaerellaceae | Mycosphaerella | *Mycosphaerella sp* | OTU 183 | 7 | 0.43 | ** | 0.01 |
| Fungi | Ascomycota | Sordariomycetes | Xylariales | Xylariaceae | Annulohypoxylon | *Annulohypoxylon urceolatum* | OTU 279 | 7 | 0.35 | ** | 0.02 |
| Fungi | Ascomycota | Dothideomycetes | Capnodiales | Mycosphaerellaceae | Zasmidium | *Zasmidium commune* | OTU 286 | 7 | 0.33 | ** | 0.04 |
| ***H. concinna*** | | | | | | | | | | | |
| Fungi | Ascomycota | Sordariomycetes | Hypocreales | Nectriaceae | Fusarium | *Fusarium sp* | OTU 7 | 5 | 0.33 | ** | 0.01 |
| Fungi | Ascomycota | Eurotiomycetes | Chaetothyriales | Herpotrichiellaceae | Exophiala | *Exophiala oligosperma* | OTU 21 | 5 | 0.22 | ** | 0.01 |
| Fungi | Ascomycota | Sordariomycetes | Hypocreales | Clavicipitaceae | unidentified | *Clavicipitaceae sp* | OTU 53 | 5 | 0.28 | ** | 0.01 |
| Fungi | Ascomycota | Saccharomycetes | Saccharomycetales | Saccharomycetales_fam_Incertae_sedis | Candida | *Candida parapsilosis* | OTU 67 | 5 | 0.27 | ** | 0.03 |
| Fungi | Ascomycota | Eurotiomycetes | Chaetothyriales | Trichomeriaceae | Bradymyces | *Bradymyces sp* | OTU 160 | 5 | 0.25 | ** | 0.05 |
| Fungi | Ascomycota | Eurotiomycetes | Chaetothyriales | Herpotrichiellaceae | Exophiala | *Exophiala oligosperma* | OTU 173 | 5 | 0.35 | ** | 0.01 |
| Fungi | Ascomycota | Sordariomycetes | Sordariales | Chaetomiaceae | unidentified | *Chaetomiaceae sp* | OTU 209 | 5 | 0.35 | ** | 0.02 |
| Fungi | Ascomycota | Dothideomycetes | Pleosporales | Didymellaceae | Neodidymelliopsis | *Neodidymelliopsis sambuci* | OTU 596 | 5 | 0.35 | ** | 0.02 |
| ***Dipteryx sp.*** | | | | | | | | | | | |
| Fungi | Ascomycota | Dothideomycetes | Capnodiales | Dissoconiaceae | Uwebraunia | *Uwebraunia dekkeri* | OTU 12 | 4 | 0.42 | ** | 0.01 |

| | | | | | | | | | | | |
|---|---|---|---|---|---|---|---|---|---|---|---|
| Fungi | Ascomycota | Eurotiomycetes | Eurotiales | Aspergillaceae | Aspergillus | *Aspergillus sp* | OTU 122 | 4 | 0.26 | ** | 0.05 |
| Fungi | Ascomycota | Dothideomycetes | Capnodiales | Dissoconiaceae | Ramichloridium | *Ramichloridium punctatum* | OTU 151 | 4 | 0.38 | ** | 0.01 |
| Fungi | Ascomycota | Sordariomycetes | Xylariales | Xylariaceae | unidentified | *Xylariaceae sp* | OTU 216 | 4 | 0.39 | ** | 0.02 |
| Fungi | Ascomycota | Dothideomycetes | Capnodiales | Schizothyriaceae | Zygophiala | *Zygophiala qianensis* | OTU 305 | 4 | 0.33 | ** | 0.03 |
| ***A. membranacea*** | | | | | | | | | | | |
| Fungi | Ascomycota | Sordariomycetes | Xylariales | unidentified | unidentified | *Xylariales sp* | OTU 25 | 1 | 0.22 | ** | 0.01 |
| Fungi | Ascomycota | Dothideomycetes | Capnodiales | Mycosphaerellaceae | Septoria | *Septoria sp* | OTU 26 | 1 | 0.37 | ** | 0.01 |
| Fungi | Ascomycota | Sordariomycetes | Xylariales | Xylariaceae | Xylaria | *Xylaria curta* | OTU 172 | 9 | 0.41 | ** | 0.01 |
| Fungi | Ascomycota | Dothideomycetes | Pleosporales | Pleosporaceae | Curvularia | *Curvularia sp* | OTU 120 | 1 | 0.35 | ** | 0.05 |
| ***C. alliodora*** | | | | | | | | | | | |
| Fungi | Ascomycota | Dothideomycetes | Botryosphaeriales | Phyllostictaceae | Phyllosticta | *Phyllosticta capitalensis* | OTU 32 | 19 | 0.36 | ** | 0.01 |
| Fungi | Ascomycota | Sordariomycetes | Diaporthales | Diaporthaceae | Diaporthe | *Diaporthe longicolla* | OTU 31 | 3 | 0.53 | ** | 0.01 |
| Fungi | Ascomycota | Sordariomycetes | Glomerellales | Glomerellaceae | Colletotrichum | *Colletotrichum gigasporum* | OTU 34 | 3 | 0.37 | ** | 0.01 |
| Fungi | Ascomycota | Sordariomycetes | Xylariales | Xylariaceae | Xylaria | *Xylaria sp* | OTU 62 | 3 | 0.52 | ** | 0.01 |
| Fungi | Ascomycota | unidentified | unidentified | unidentified | unidentified | *Ascomycota sp* | OTU 52 | 3 | 0.49 | ** | 0.01 |
| Fungi | Ascomycota | Sordariomycetes | Diaporthales | Diaporthaceae | Diaporthe | *Diaporthe sp* | OTU 99 | 3 | 0.44 | ** | 0.01 |
| Fungi | Ascomycota | Sordariomycetes | Sordariales | Chaetomiaceae | Ovatospora | *Ovatospora brasiliensis* | OTU 61 | 3 | 0.37 | ** | 0.01 |
| Fungi | Ascomycota | Dothideomycetes | Pleosporales | Phaeosphaeriaceae | Setophoma | *Setophoma sp* | OTU 71 | 3 | 0.39 | ** | 0.01 |
| Fungi | Ascomycota | Sordariomycetes | Xylariales | Xylariaceae | Annulohypoxylon | *Annulohypoxylon stygium* | OTU 78 | 3 | 0.5 | ** | 0.01 |
| Fungi | Ascomycota | Sordariomycetes | Xylariales | Xylariaceae | Hypoxylon | *Hypoxylon sp* | OTU 101 | 3 | 0.48 | ** | 0.01 |
| Fungi | Ascomycota | Sordariomycetes | Diaporthales | Diaporthaceae | Diaporthe | *Diaporthe sp* | OTU 223 | 3 | 0.42 | ** | 0.01 |
| Fungi | Ascomycota | Sordariomycetes | Xylariales | Xylariaceae | Xylaria | *Xylaria sp* | OTU 212 | 3 | 0.5 | ** | 0.01 |
| Fungi | Ascomycota | Dothideomycetes | Capnodiales | Cladosporiaceae | Melomastia | *Melomastia sp* | OTU 117 | 3 | 0.43 | ** | 0.01 |
| Fungi | Ascomycota | Dothideomycetes | Pleosporales | Morosphaeriaceae | Acrocalymma | *Acrocalymma sp* | OTU 169 | 3 | 0.43 | ** | 0.01 |
| Fungi | Ascomycota | Sordariomycetes | Diaporthales | Diaporthaceae | Diaporthe | *Diaporthe sp* | OTU 512 | 3 | 0.46 | ** | 0.01 |
| Fungi | Ascomycota | Sordariomycetes | Diaporthales | Diaporthaceae | Diaporthe | *Diaporthe melonis* | OTU 179 | 3 | 0.48 | ** | 0.01 |
| Fungi | Ascomycota | Sordariomycetes | Xylariales | Xylariales_fam_Incertae_sedis | Oxydothis | *Oxydothis garethjonesii* | OTU 94 | 57 | 0.37 | ** | 0.03 |
| Fungi | Ascomycota | Sordariomycetes | Xylariales | Xylariaceae | unidentified | *Xylariaceae sp* | OTU 106 | 3 | 0.46 | ** | 0.01 |

| | | | | | | | | | | | |
|---|---|---|---|---|---|---|---|---|---|---|---|
| Fungi | Ascomycota | Sordariomycetes | Xylariales | Xylariaceae | Hypoxylon | *Hypoxylon hypomiltum* | OTU 142 | 3 | 0.37 | ** | 0.01 |
| Fungi | Ascomycota | Sordariomycetes | Xylariales | Xylariaceae | Xylaria | *Xylaria sp* | OTU 262 | 3 | 0.4 | ** | 0.01 |
| Fungi | Ascomycota | Sordariomycetes | Diaporthales | Diaporthaceae | Diaporthe | *Diaporthe sp* | OTU 327 | 3 | 0.48 | ** | 0.01 |
| Fungi | Ascomycota | Sordariomycetes | Xylariales | Xylariaceae | Hypoxylon | *Hypoxylon submonticulosum* | OTU 202 | 3 | 0.34 | ** | 0.01 |
| Fungi | Ascomycota | Sordariomycetes | Xylariales | Amphisphaeriaceae | Lepteutypa | *Lepteutypa sambuci* | OTU 210 | 3 | 0.41 | ** | 0.01 |
| Fungi | Ascomycota | Sordariomycetes | Xylariales | Xylariaceae | unidentified | *Xylariaceae sp* | OTU 380 | 3 | 0.35 | ** | 0.02 |
| Fungi | Ascomycota | Sordariomycetes | Xylariales | Xylariaceae | unidentified | *Xylariaceae sp* | OTU 537 | 3 | 0.45 | ** | 0.01 |
| Fungi | Ascomycota | Sordariomycetes | Sordariales | Lasiosphaeriaceae | unidentified | *Lasiosphaeriaceae sp* | OTU 219 | 3 | 0.35 | ** | 0.02 |
| Fungi | Ascomycota | unidentified | unidentified | unidentified | unidentified | *Ascomycota sp* | OTU 192 | 3 | 0.4 | ** | 0.01 |
| Fungi | Ascomycota | Sordariomycetes | Xylariales | Xylariaceae | Lopadostoma | *Lopadostoma americanum* | OTU 936 | 3 | 0.42 | ** | 0.02 |
| Fungi | Ascomycota | Sordariomycetes | Sordariales | Sordariales_fam_Incertae_sedis | Ramophialophora | *Ramophialophora sp* | OTU 362 | 3 | 0.33 | ** | 0.04 |
| Fungi | Ascomycota | unidentified | unidentified | unidentified | unidentified | *Ascomycota sp* | OTU 784 | 3 | 0.4 | ** | 0.01 |
| Fungi | Ascomycota | Sordariomycetes | unidentified | unidentified | unidentified | *Sordariomycetes sp* | OTU 397 | 3 | 0.44 | ** | 0.02 |
| Fungi | Ascomycota | Sordariomycetes | Diaporthales | Diaporthaceae | Diaporthe | *Diaporthe fraxini-angustifoliae* | OTU 575 | 3 | 0.38 | ** | 0.02 |
| Fungi | Ascomycota | Sordariomycetes | Glomerellales | Glomerellaceae | Colletotrichum | *Colletotrichum ignotum* | OTU 614 | 3 | 0.34 | ** | 0.03 |
| ***C. cainito*** | | | | | | | | | | | |
| Fungi | Ascomycota | Sordariomycetes | Xylariales | unidentified | unidentified | *Xylariales sp* | OTU 58 | 2 | 0.36 | ** | 0.03 |
| Fungi | Ascomycota | Sordariomycetes | Hypocreales | Hypocreales_fam_Incertae_sedis | Sarocladium | *Sarocladium gamsii* | OTU 75 | 18 | 0.34 | ** | 0.01 |
| Fungi | Ascomycota | Sordariomycetes | Diaporthales | Valsaceae | Phomopsis | *Phomopsis sp* | OTU 86 | 2 | 0.32 | ** | 0.01 |
| Fungi | Ascomycota | Sordariomycetes | Sordariales | Cephalothecaceae | Phialemonium | *Phialemonium dimorphosporum* | OTU 139 | 2 | 0.33 | ** | 0.02 |
| Fungi | Ascomycota | Sordariomycetes | Hypocreales | Hypocreales_fam_Incertae_sedis | Acremonium | *Acremonium hennebertii* | OTU 145 | 2 | 0.41 | ** | 0.01 |
| Fungi | Ascomycota | Sordariomycetes | Xylariales | unidentified | unidentified | *Xylariales sp* | OTU 128 | 50 | 0.34 | ** | 0.04 |
| Fungi | Ascomycota | Arthoniomycetes | Lichenostigmatales | Phaeococcomycetaceae | Phaeococcomyces | *Phaeococcomyces rothmanniae* | OTU 197 | 2 | 0.35 | ** | 0.01 |
| Fungi | Ascomycota | Sordariomycetes | Diaporthales | Diaporthaceae | Diaporthe | *Diaporthe sp* | OTU 761 | 2 | 0.42 | ** | 0.01 |

| | | | | | | | | | | |
|---|---|---|---|---|---|---|---|---|---|---|
| ***L. panamensis*** | | | | | | | | | | |
| Fungi | Ascomycota | Sordariomycetes | Xylariales | Xylariales_fam_Incertae_sedis | Oxydothis | *Oxydothis sp* | OTU 221 | 6 | 0.34 | ** | 0.03 |
| Fungi | Ascomycota | Dothideomycetes | Pleosporales | unidentified | unidentified | *Pleosporales sp* | OTU 232 | 6 | 0.34 | ** | 0.03 |
| Fungi | Ascomycota | Sordariomycetes | Glomerellales | Glomerellales_fam_Incertae_sedis | Malaysiasca | *Malaysiasca phaii* | OTU 735 | 6 | 0.31 | ** | 0.02 |

[1] Significance levels are represented by *ast*erisks [*p* < .05 (*), *p* <= .01 (**), *p* <= .001 (***), and *p* < .0001 (****)]..

[2]Benjamini & Hochberg method adjustment for multiple comparisons

#### Table S6: Taxonomy of significantly correlated OTUs with *Atta colombica* herbivory levels

| | | | | | | | | Multilevel pattern analysis | | | |
|---|---|---|---|---|---|---|---|---|---|---|---|
| **Kingdom** | **Phylum** | **Class** | **Order** | **Family** | **Genus** | **Species** | **OTU** | **Index** | **Stat** | ***p*[2]** | ***p*adj[3]** |
| **Medium** | | | | | | | | | | | |
| Fungi | Ascomycota | Eurotiomycetes | Chaetothyriales | Cyphellophoraceae | Cyphellophora | *Cyphellophora oxyspora* | OTU 19 | 3 | 0.29 | * | 1 |
| Fungi | Ascomycota | unidentified | unidentified | unidentified | unidentified | *Ascomycota sp* | OTU 153 | 3 | 0.31 | * | 1 |
| Fungi | Ascomycota | Sordariomycetes | Hypocreales | unidentified | unidentified | *Hypocreales sp* | OTU 682 | 3 | 0.31 | * | 1 |
| **High** | | | | | | | | | | | |
| Fungi | Ascomycota | Sordariomycetes | Xylariales | Xylariaceae | Xylaria | *Xylaria sp* | OTU 62 | 1 | 0.32 | * | 1 |
| Fungi | Ascomycota | Eurotiomycetes | Eurotiales | Aspergillaceae | Aspergillus | *Aspergillus terreus* | OTU 55 | 1 | 0.23 | * | 1 |
| Fungi | Ascomycota | Sordariomycetes | Xylariales | Xylariaceae | Xylaria | *Xylaria sp* | OTU 212 | 1 | 0.34 | * | 1 |
| Fungi | Ascomycota | Sordariomycetes | Xylariales | Xylariaceae | unidentified | *Xylariaceae sp* | OTU 106 | 1 | 0.31 | * | 1 |
| Fungi | Ascomycota | Sordariomycetes | Microascales | unidentified | unidentified | *Microascales sp* | OTU 608 | 1 | 0.27 | * | 1 |
| Fungi | Ascomycota | Sordariomycetes | Hypocreales | Cordycipitaceae | Beauveria | *Beauveria sp* | OTU 204 | 1 | 0.28 | * | 1 |
| Fungi | Ascomycota | unidentified | unidentified | unidentified | unidentified | *Ascomycota sp* | OTU 784 | 1 | 0.26 | * | 1 |
| Fungi | Ascomycota | Sordariomycetes | Sordariales | unidentified | unidentified | *Sordariales sp* | OTU 437 | 1 | 0.3 | * | 1 |
| Fungi | Ascomycota | Sordariomycetes | unidentified | unidentified | unidentified | *Sordariomycetes sp* | OTU 492 | 1 | 0.26 | * | 1 |
| Fungi | Ascomycota | Dothideomycetes | Pleosporales | Lentitheciaceae | Poaceascoma | *Poaceascoma sp* | OTU 644 | 1 | 0.28 | * | 1 |
| Fungi | Ascomycota | Sordariomycetes | Microascales | unidentified | unidentified | *Microascales sp* | OTU 1043 | 1 | 0.3 | * | 1 |
| Fungi | Ascomycota | Sordariomycetes | Microascales | Halosphaeriaceae | unidentified | *Halosphaeriaceae sp* | OTU 1053 | 1 | 0.25 | * | 1 |
| Fungi | Ascomycota | Sordariomycetes | Xylariales | Xylariales_fam_Incertae_sedis | Phialemoniopsis | *Phialemoniopsis sp* | OTU 1067 | 1 | 0.3 | * | 1 |
| **Low** | | | | | | | | | | | |
| Fungi | Ascomycota | Sordariomycetes | Hypocreales | Hypocreales_fam_Incertae_sedis | Acremonium | *Acremonium hennebertii* | OTU 100 | 2 | 0.32 | ** | 1 |

[1]High = >70% leaf area damage, Medium = 31-69% leaf area damage, Low = <30% leaf area damage

[2]Significance levels are represented by asterisks [$p < .05$ (*), $p <= .01$ (**), $p <= .001$ (***), and $p < .0001$ (****)].

[3]Benjamini & Hochberg method adjustment for multiple comparisons

#### Table S7: Taxonomy of significantly correlated OTUs with *Calonectria* sp. pathogen damage levels

| | | | | | | | | Multilevel pattern analysis | | | |
|---|---|---|---|---|---|---|---|---|---|---|---|
| **Kingdom** | **Phylum** | **Class** | **Order** | **Family** | **Genus** | **Species** | **OTU** | **Index** | **Stat** | ***p*[2]** | ***p*adj[3]** |
| **High** | | | | | | | | | | | |
| Fungi | Ascomycota | Sordariomycetes | Glomerellales | Glomerellaceae | Colletotrichum | *Colletotrichum fructicola* | OTU 1 | 1 | 0.22 | * | 1 |
| Fungi | Ascomycota | Dothideomycetes | Capnodiales | Dissoconiaceae | Ramichloridium | *Ramichloridium apiculatum* | OTU 18 | 1 | 0.24 | * | 1 |
| Fungi | Ascomycota | Dothideomycetes | Botryosphaeriales | Phyllostictaceae | Phyllosticta | *Phyllosticta capitalensis* | OTU 32 | 1 | 0.23 | ns | 1 |
| Fungi | Ascomycota | Dothideomycetes | Pleosporales | unidentified | unidentified | *Pleosporales sp* | OTU 70 | 1 | 0.23 | * | 1 |
| Fungi | Ascomycota | Sordariomycetes | unidentified | unidentified | unidentified | *Sordariomycetes sp* | OTU 114 | 1 | 0.19 | * | 1 |
| Fungi | Ascomycota | Dothideomycetes | Pleosporales | Leptosphaeriaceae | Leptosphaeria | *Leptosphaeria modesta* | OTU 108 | 1 | 0.24 | * | 1 |
| Fungi | Ascomycota | Sordariomycetes | Chaetosphaeriales | Chaetosphaeriaceae | unidentified | *Chaetosphaeriaceae sp* | OTU 177 | 1 | 0.2 | ns | 1 |
| Fungi | Ascomycota | Sordariomycetes | Xylariales | Xylariaceae | Hypoxylon | *Hypoxylon sp* | OTU 244 | 1 | 0.21 | * | 1 |
| Fungi | Ascomycota | Sordariomycetes | Hypocreales | unidentified | unidentified | *Hypocreales sp* | OTU 196 | 1 | 0.23 | ns | 1 |
| Fungi | Ascomycota | Sordariomycetes | Sordariales | Lasiosphaeriaceae | unidentified | *Lasiosphaeriaceae sp* | OTU 258 | 1 | 0.23 | * | 1 |
| Fungi | Ascomycota | Sordariomycetes | Phomatosporales | Phomatosporaceae | Phomatospora | *Phomatospora sp* | OTU 637 | 1 | 0.21 | * | 1 |

[1]High = >30% leaf area damage, Low = <29% leaf area damage

[2]Significance levels are represented by *ast*erisks [*p* < .05 (*), *p* <= .01 (**), *p* <= .001 (***), and *p* < .0001 (****)].

[3]Benjamini & Hochberg method adjustment for multiple comparisons

#### Table S8: Taxonomy of significantly correlated OTUs with FEF inoculation levels

| | | | | | | | | Multilevel pattern analysis | | | |
|---|---|---|---|---|---|---|---|---|---|---|---|
| **Kingdom** | **Phylum** | **Class** | **Order** | **Family** | **Genus** | **Species** | **OTU** | **Index** | **Stat** | ***p*[1]** | ***p*adj[2]** |
| **E+** | | | | | | | | | | | |
| Fungi | Ascomycota | Sordariomycetes | Glomerellales | Glomerellaceae | Colletotrichum | *Colletotrichum fructicola* | OTU 1 | 2 | 0.11 | ** | 0.02 |
| Fungi | Ascomycota | Sordariomycetes | Xylariales | Sporocadaceae | Neopestalotiopsis | *Neopestalotiopsis sp* | OTU 10 | 2 | 0.2 | ** | 0.02 |
| Fungi | Ascomycota | Dothideomycetes | Capnodiales | Dissoconiaceae | Uwebraunia | *Uwebraunia dekkeri* | OTU 12 | 2 | 0.2 | ** | 0.02 |
| Fungi | Ascomycota | Sordariomycetes | unidentified | unidentified | unidentified | *Sordariomycetes sp* | OTU 36 | 2 | 0.42 | ** | 0.02 |
| Fungi | Ascomycota | Sordariomycetes | Hypocreales | unidentified | unidentified | *Hypocreales sp* | OTU 39 | 2 | 0.3 | ** | 0.02 |
| Fungi | Ascomycota | Sordariomycetes | Diaporthales | Diaporthaceae | Diaporthe | *Diaporthe longicolla* | OTU 31 | 2 | 0.21 | ** | 0.02 |
| Fungi | Ascomycota | Sordariomycetes | Glomerellales | Plectosphaerellaceae | Wallrothiella | *Wallrothiella subiculosa* | OTU 35 | 2 | 0.27 | ** | 0.02 |
| Fungi | Ascomycota | Sordariomycetes | Glomerellales | Glomerellaceae | Colletotrichum | *Colletotrichum gigasporum* | OTU 34 | 2 | 0.27 | ** | 0.02 |
| Fungi | Ascomycota | Sordariomycetes | unidentified | unidentified | unidentified | *Sordariomycetes sp* | OTU 42 | 2 | 0.42 | ** | 0.02 |
| Fungi | Ascomycota | Sordariomycetes | Xylariales | Sporocadaceae | Pseudopestalotiopsis | *Pseudopestalotiopsis sp* | OTU 46 | 2 | 0.19 | ** | 0.02 |
| Fungi | Ascomycota | Sordariomycetes | Hypoceales | Amplistromataceae | Amplistroma | *Amplistroma erinaceum* | OTU 60 | 2 | 0.32 | ** | 0.02 |
| Fungi | Ascomycota | Sordariomycetes | Xylariales | Xylariaceae | Xylaria | *Xylaria sp* | OTU 62 | 2 | 0.24 | ** | 0.02 |
| Fungi | Ascomycota | Sordariomycetes | Hypocreales | Hypocreales_fam_Incertae_sedis | Acremonium | *Acremonium sp* | OTU 49 | 2 | 0.17 | ** | 0.02 |
| Fungi | Ascomycota | unidentified | unidentified | unidentified | unidentified | *Ascomycota sp* | OTU 52 | 2 | 0.29 | ** | 0.02 |
| Fungi | Ascomycota | Sordariomycetes | Diaporthales | Diaporthaceae | Diaporthe | *Diaporthe sp* | OTU 99 | 2 | 0.21 | ** | 0.04 |
| Fungi | Ascomycota | Dothideomycetes | Capnodiales | Cladosporiaceae | Cladosporium | *Cladosporium sp* | OTU 132 | 2 | 0.2 | ** | 0.04 |
| Fungi | Ascomycota | Sordariomycetes | Hypocreales | Hypocreales_fam_Incertae_sedis | Acremonium | *Acremonium hennebertii* | OTU 77 | 2 | 0.27 | ** | 0.02 |
| Fungi | Ascomycota | Sordariomycetes | Diaporthales | Diaporthaceae | Diaporthe | *Diaporthe sp* | OTU 223 | 2 | 0.18 | ** | 0.02 |
| Fungi | Ascomycota | Sordariomycetes | Hypocreales | Hypocreales_fam_Incertae_sedis | Acremonium | *Acremonium hennebertii* | OTU 100 | 2 | 0.26 | ** | 0.02 |
| Fungi | Ascomycota | Dothideomycetes | Capnodiales | Cladosporiaceae | Melomastia | *Melomastia sp* | OTU 117 | 2 | 0.17 | ** | 0.04 |
| Fungi | Ascomycota | Sordariomycetes | unidentified | unidentified | unidentified | *Sordariomycetes sp* | OTU 92 | 2 | 0.29 | ** | 0.02 |
| Fungi | Ascomycota | Dothideomy | Pleosporale | Leptosphaer | Leptosphaer | *Leptosphae* | OTU 108 | 2 | 0.28 | ** | 0.02 |

| | | | | | | | | | | | |
|---|---|---|---|---|---|---|---|---|---|---|---|
| | | cetes | s | iaceae | ia | *ria modesta* | | | | | |
| Fungi | Ascomycota | Sordariomycetes | Diaporthales | Diaporthaceae | Diaporthe | *Diaporthe melonis* | OTU 179 | 2 | 0.22 | ** | 0.04 |
| Fungi | Ascomycota | Sordariomycetes | Xylariales | Xylariales_fam_Incertae_sedis | Oxydothis | *Oxydothis garethjonesii* | OTU 94 | 2 | 0.3 | ** | 0.02 |
| Fungi | Ascomycota | Sordariomycetes | Xylariales | Xylariaceae | unidentified | *Xylariaceae sp* | OTU 106 | 2 | 0.26 | ** | 0.02 |
| Fungi | Ascomycota | Eurotiomycetes | Chaetothyriales | unidentified | unidentified | *Chaetothyriales sp* | OTU 126 | 2 | 0.31 | ** | 0.02 |
| Fungi | Ascomycota | Sordariomycetes | Glomerellales | Plectosphaerellaceae | unidentified | *Plectosphaerellaceae sp* | OTU 146 | 2 | 0.13 | ** | 0.04 |
| Fungi | Ascomycota | Sordariomycetes | Sordariomycetes_ord_Incertae_sedis | Sordariomycetes_fam_Incertae_sedis | Distoseptispora | *Distoseptispora sp* | OTU 148 | 2 | 0.29 | ** | 0.02 |
| Fungi | Ascomycota | Eurotiomycetes | Chaetothyriales | Herpotrichiellaceae | Phialophora | *Phialophora geniculata* | OTU 278 | 2 | 0.3 | ** | 0.02 |
| Fungi | Ascomycota | Sordariomycetes | Glomerellales | Plectosphaerellaceae | Plectosphaerella | *Plectosphaerella cucumerina* | OTU 390 | 2 | 0.2 | ** | 0.02 |
| Fungi | Ascomycota | Dothideomycetes | unidentified | unidentified | unidentified | *Dothideomycetes sp* | OTU 201 | 2 | 0.27 | ** | 0.02 |

[1]Significance levels are represented by *ast*erisks [*p* < .05 (*), *p* <= .01 (**), *p* <= .001 (***), and *p* < .0001 (****)].

[2]Benjamini & Hochberg method adjustment for multiple comparisons

**Table S9: Sterilization protocol for tropical tree seeds**

| Tree Species | Number of seed collected | Number of maternal sources | Sterilization protocol |
|---|---|---|---|
| *Apeiba membranacea* | 500 | 3 | Soak in water 3-5 days; 0.5% NaClO for 4 minutes; 70% EtOH for 5 minutes |
| *Chrysophylum cainito* | 100 | 1 | Soak in water 3-5 days; 0.5% NaClO for 4 minutes, 70% EtOH for 5 minutes |
| *Cordia alliodora* | 403 | 1 | Soak in water 1 day; 0.5% NaClo for 3 minutes; 50% EtOH for 3 minutes |
| *Dipteryx sp.* | ~100 | 1 | Soak in water 7 days; 0.5% NaClO for 5 minutes; 70% EtOH for 5 minutes |
| *Heisteria concinna* | 250 | ~6 | Soak in water 3-5 days; 0.5% NaClO for 4 minutes; 70% EtOH for 5 minutes |
| *Lacmellea panamensis* | 75 | 3 | Soak in water 3-5 days; 0.25% NaClO for 3 minutes; 50% EtOH for 3 minutes |
| *Theobroma cacao* | 44 | 1 | Rinsed seeds in running tap water; 0.5% NaClO for 5 minutes |

**Figure S1**

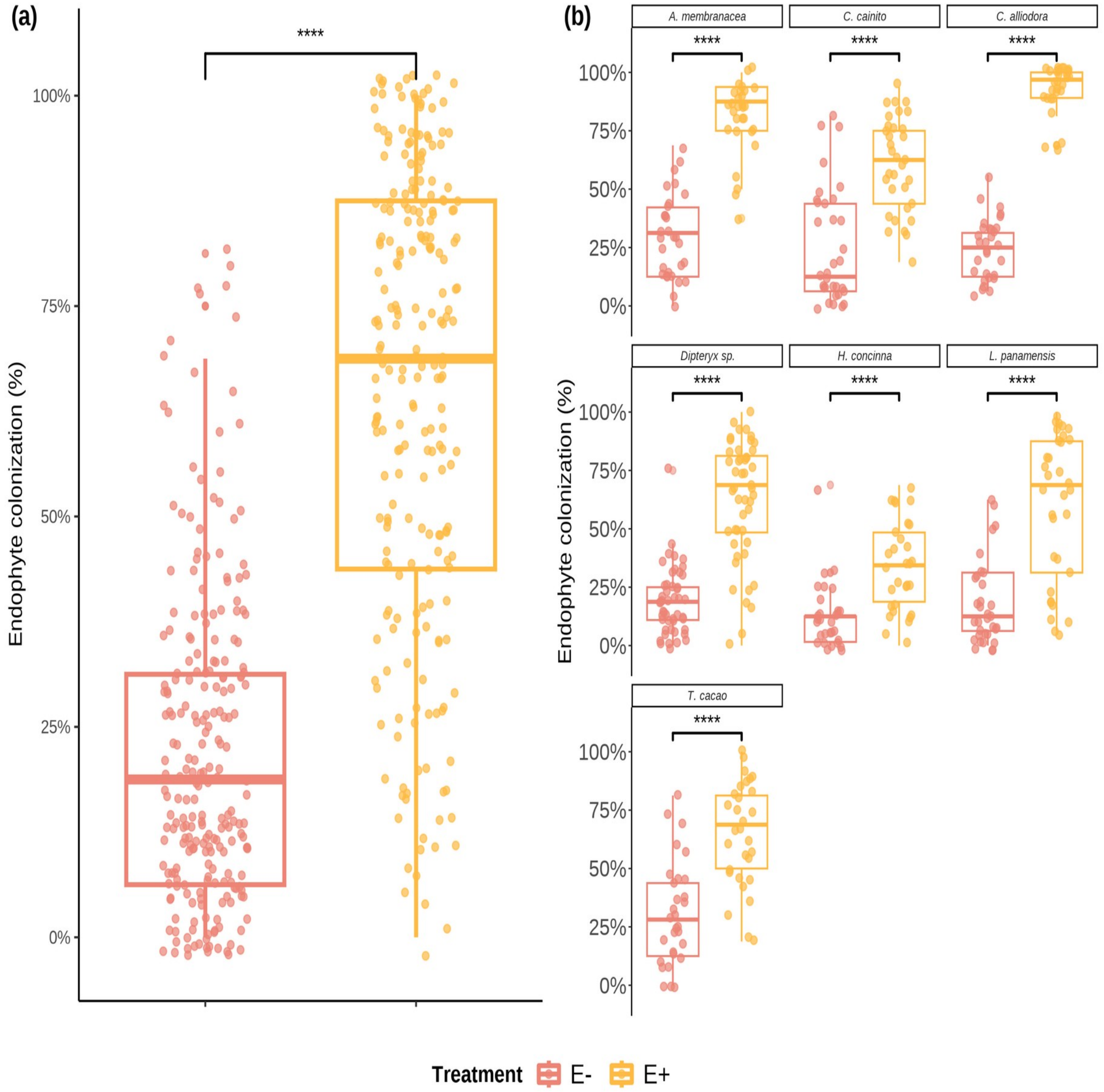


*Foliar endophytic fungi (FEF) colonization of seven tropical tree species in malt extract agar (MEA 2%). a) Comparison of mean percent colonization of leaves by FEF measured 7 days after placing leaf pieces on plates. Statistical significance was calculated with a Student's t-Test. Violin plots show the distribution of colonization values for all tree species within treatment groups (E- and E+). b) Comparison of mean percent colonization of leaves by FEF measured 7 days after culture. Violin plots show the distribution of percent colonization values for each species per treatment group. Pink filled violins represent low FEF group (E-) and yellow filled violins represent high FEF group (E+). Significance levels are represented by ns (not significant) and asterisks* [*p* < .05 (*), *p* < .01 (**), *p* < .001 (***), and *p* < .0001 (****)].

**Figure S2**

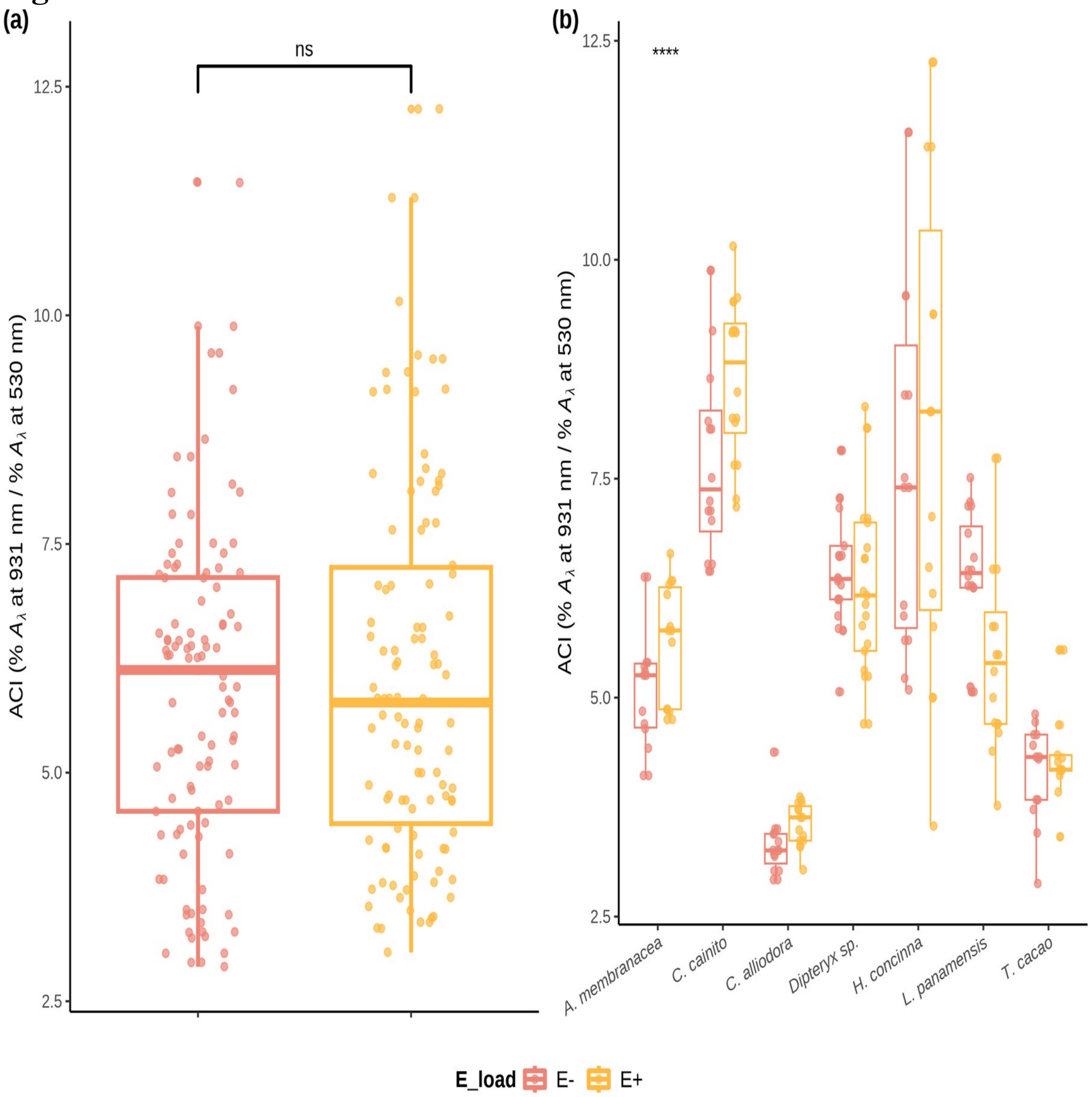


*Distributions of values and means of anthocyanin content (ACI) in treatment groups (E- and E+) and tree species. a) Comparison of ACI means between treatment groups across individuals of all species. Statistical significance was calculated using a two-sided Student's t-Test. b) Comparison of ACI means between treatment types of each species. Statistical significance was calculated with an analysis of variance (ANOVA). Pink filled violins represent low FEF group (E-) and yellow filed violins represent high FEF group (E+). Significance levels are represented by ns (not significant) and asterisks* [$p < .05$ (*), $p < .01$ (**), $p < .001$ (***), and $p < .0001$ (****)].

#### Figure S3

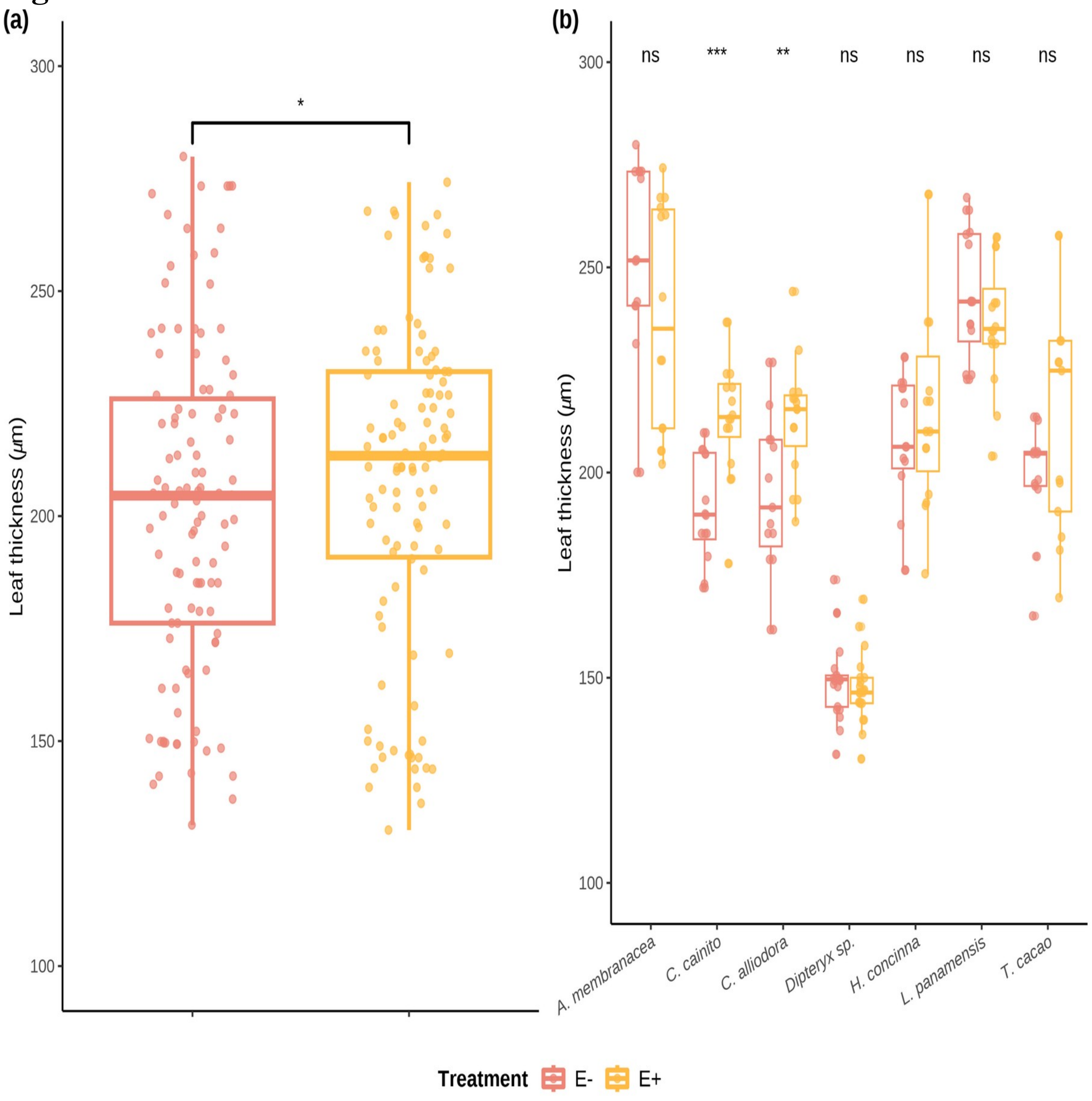


*Distributions of values and means of leaf thickness (LT) (µg) in treatment groups (E- and E+) and tree species. a) Comparison of LT means between treatment groups across individuals of all species. Statistical significance was calculated using a two-sided Student's t-Test. b) Comparison of LT means between treatment types of each species. Statistical significance was calculated with an analysis of variance (ANOVA). Pink filled violins represent low FEF group (E-) and yellow filed violins represent high FEF group (E+). Significance levels are represented by ns (not significant) and asterisks* [$p < .05$ (*), $p < .01$ (**), $p < .001$ (***), and $p < .0001$ (****)].

#### Figure S4

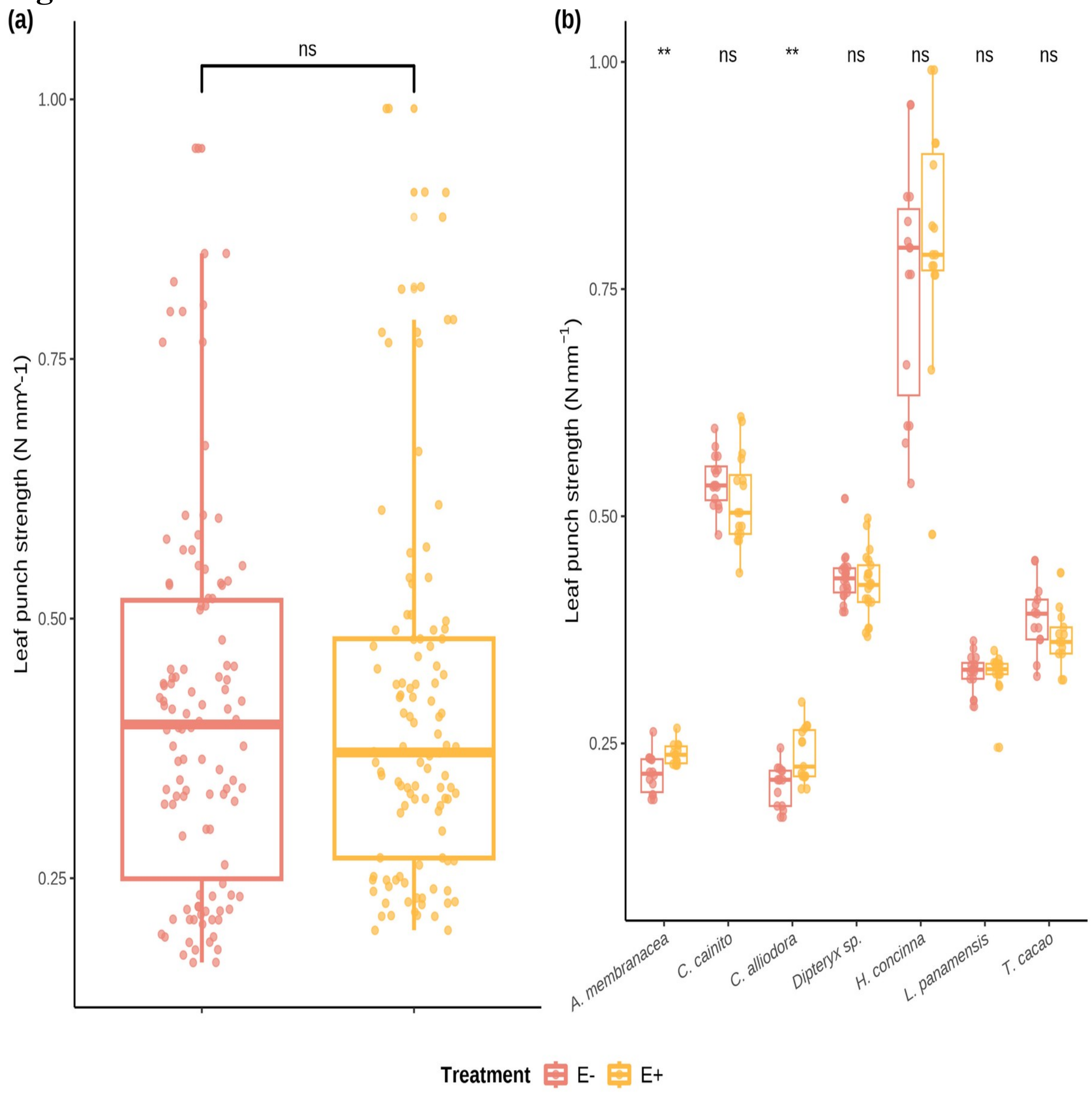


*Distributions of values and means of leaf punch strength (LPS) (N mm^-1) in treatment groups (E- and E+) and tree species. a) Comparison of LPS means between treatment groups across individuals of all species. Statistical significance was calculated using a two-sided Student's t-Test. b) Comparison of LPS means between treatment types of each species. Statistical significance was calculated with an analysis of variance (ANOVA). Pink filled violins represent low FEF group (E-) and yellow filed violins represent high FEF group (E+). Significance levels are represented by ns (not significant) and asterisks* [$p < .05$ (*), $p < .01$ (**), $p < .001$ (***), and $p < .0001$ (****)].

**Figure S5**

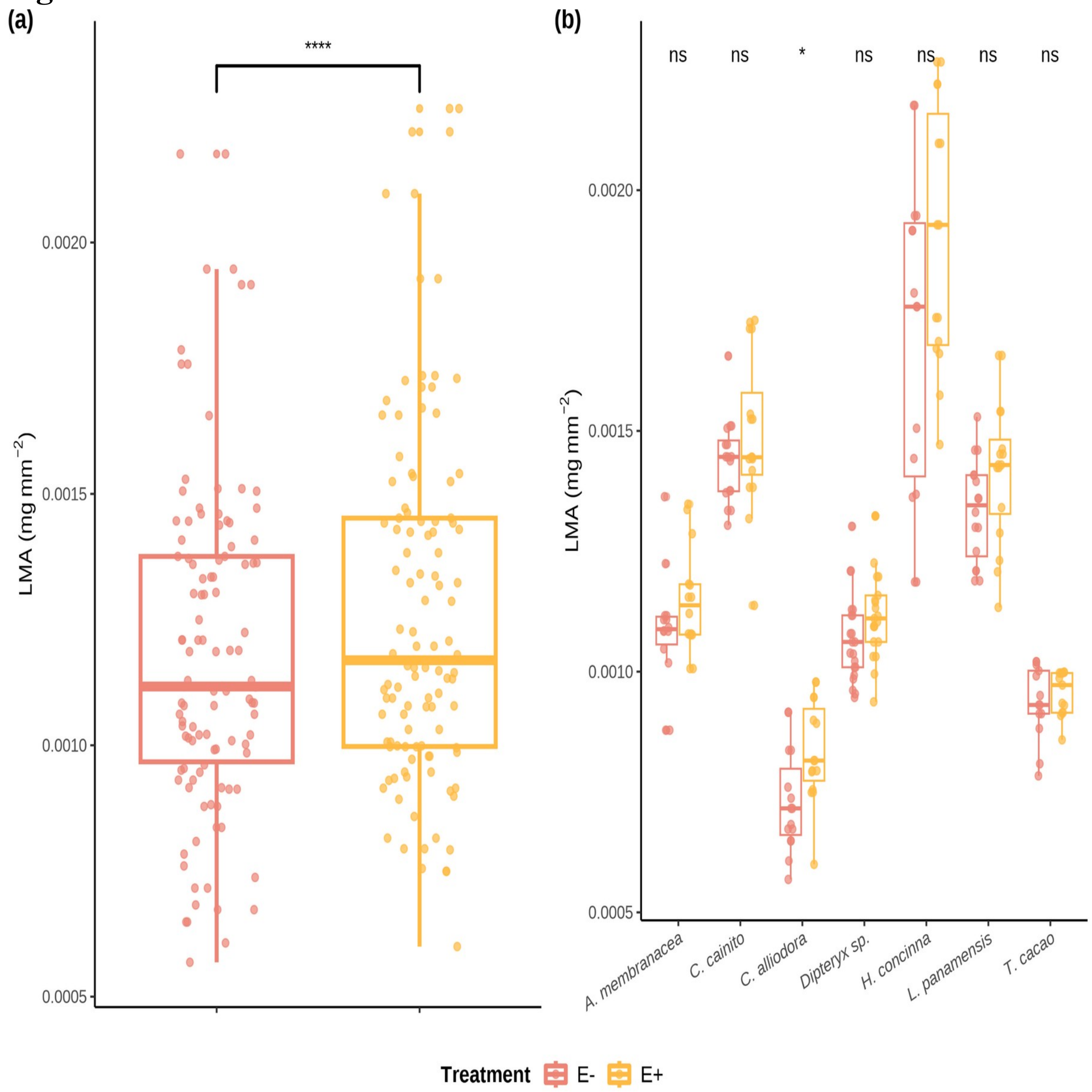


*Distributions of values and means of leaf mass per area (LMA) (mg mm^2) in treatment groups (E- and E+) and tree species. a) Comparison of LMA means between treatment groups across individuals of all species. Statistical significance was calculated using a two-sided Student's t-Test. b) Comparison of LMA means between treatment types of each species. Statistical significance was calculated with an analysis of variance (ANOVA). Pink filled violins represent low FEF group (E-) and yellow filed violins represent high FEF group (E+). Significance levels are represented by ns (not significant) and asterisks* [*p* < .05 (*), *p* < .01 (**), *p* < .001 (***), and *p* < .0001 (****)].

#### Figure S6

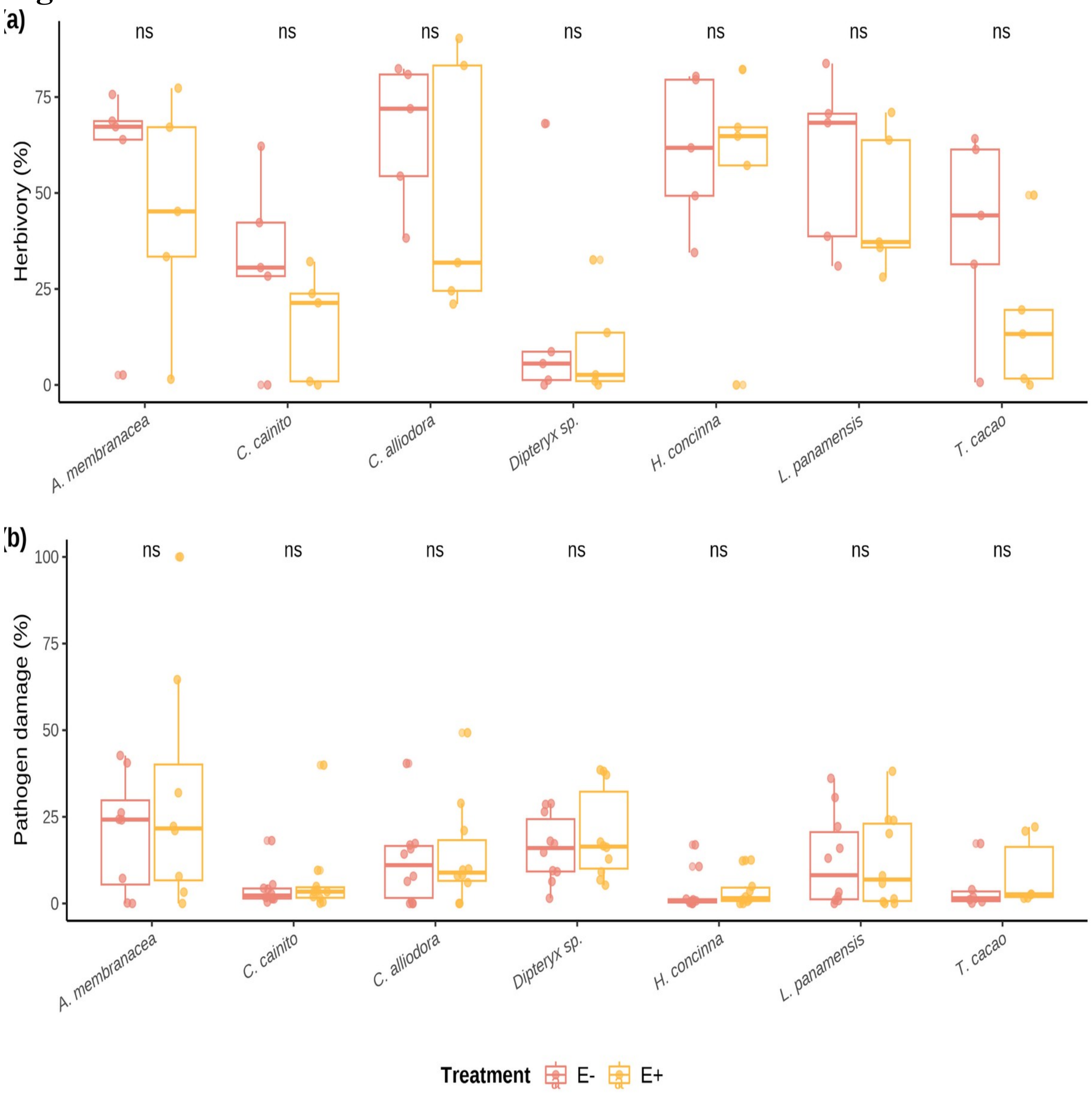


*Distributions of values and means of herbivory (%) and pathogen damage caused by Atta colombica and Calonectria sp. , respectively, in treatment groups (E- and E+) per tree species. a) Comparison of herbivory (%) means between treatment groups across individuals of all species. b) Comparison of pathogen (%) means between treatment groups across individuals of all species. Pink filled violins represent low FEF group (E-) and yellow filed violins represent high FEF group (E+). Statistical significance was calculated with an analysis of variance (ANOVA). Significance levels are represented by ns (not significant) and asterisks Significance levels are represented by ns (not significant) and asterisks* [$p < .05$ (*), $p < .01$ (**), $p < .001$ (***), and $p < .0001$ (****)].

#### Figure S7

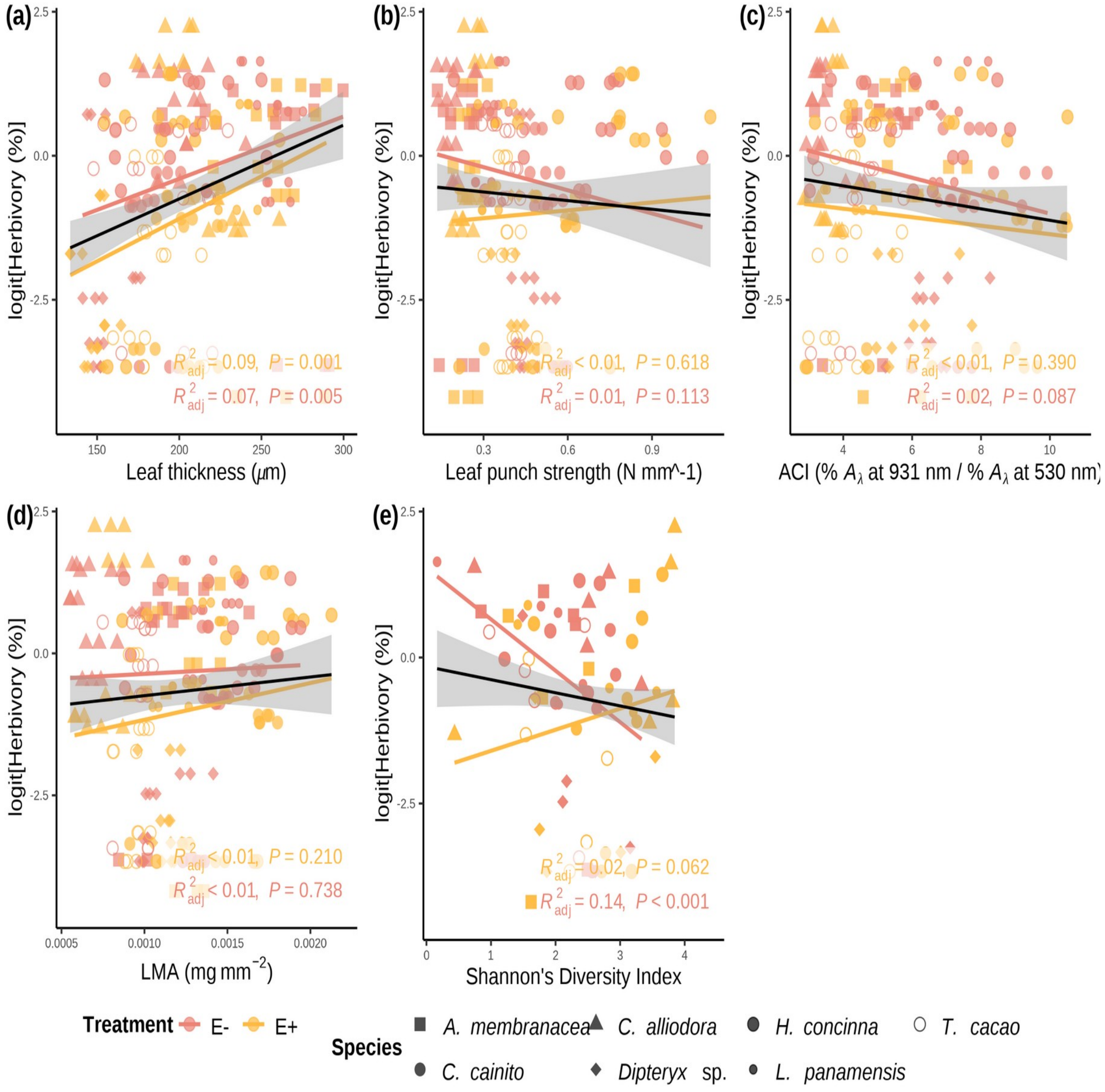


*Simple linear regressions of logit transfomed herbivory (%) and and leaf functional traits. a) Herbivory vs. leaf thickness (LT) (µg) ($R^2$-adjusted= 0.0811, p < .0001). b) Herbivory vs. LPS (N mm-1) ($R^2$-adjusted= -0.0018, p = 0.429)). c) Herbivory vs. ACI ($R^2$-adjusted= 0.0071, p = 0.116). d) Pathogen vs. LMA (mg mm^2) ($R^2$-adjusted= -0.0008, p = 0.36). e) Herbivory vs. Shannon diversity index ($R^2$-adjusted= 0.007 , p = 0.12). Pink filled line and shapes represent low FEF group (E-) and yellow filled line and shapes represent high FEF group (E+). Black line represents the linear regression on all observations.*

#### Figure S8

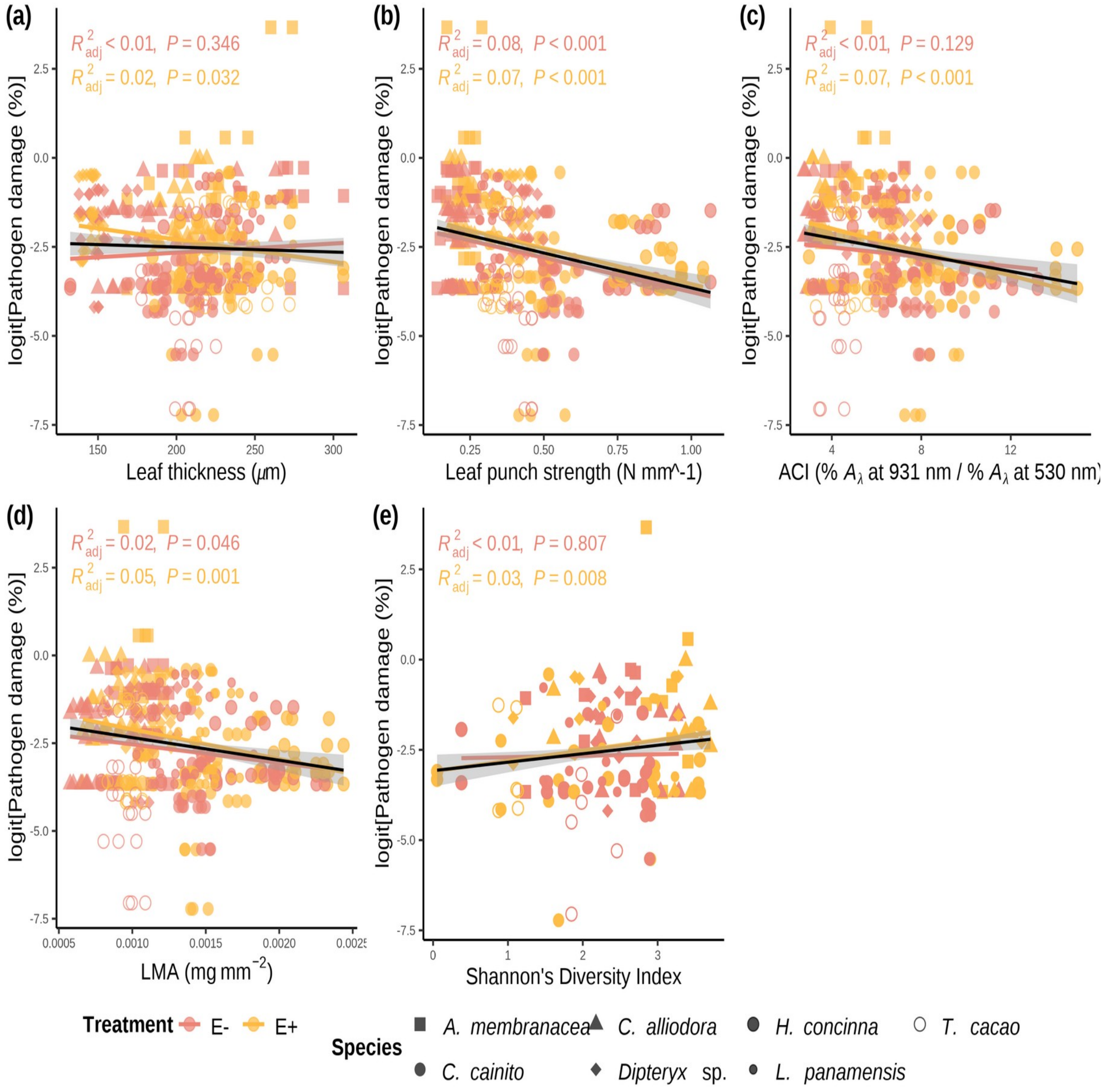


Simple linear regressions of logit transformed pathogen damage (%) and and leaf functional traits. a) Pathogen damage vs. leaf thickness (LT) (µg) ($R^2$-adjusted= -0.0013, $p$ = 0.482). b) Pathogen damage vs. LPS (N mm-1) ($R^2$-adjusted= 0.0782, $p$ < .0001)). c) Pathogen damage vs. ACI ($R^2$-adjusted= 0.0338, $p$ < .001). d) Pathogen vs. LMA (mg mm^2) ($R^2$-adjusted= 0.0295, $p$ < .001). e) Pathogen damage vs. Shannon diversity index ($R^2$-adjusted= 0.0152 , $p$ < .001). Pink filled line and shapes represent low FEF group (E-) and yellow filled line and shapes represent high FEF group (E+). Black line represents the linear regression on all observations.

### Figure S9

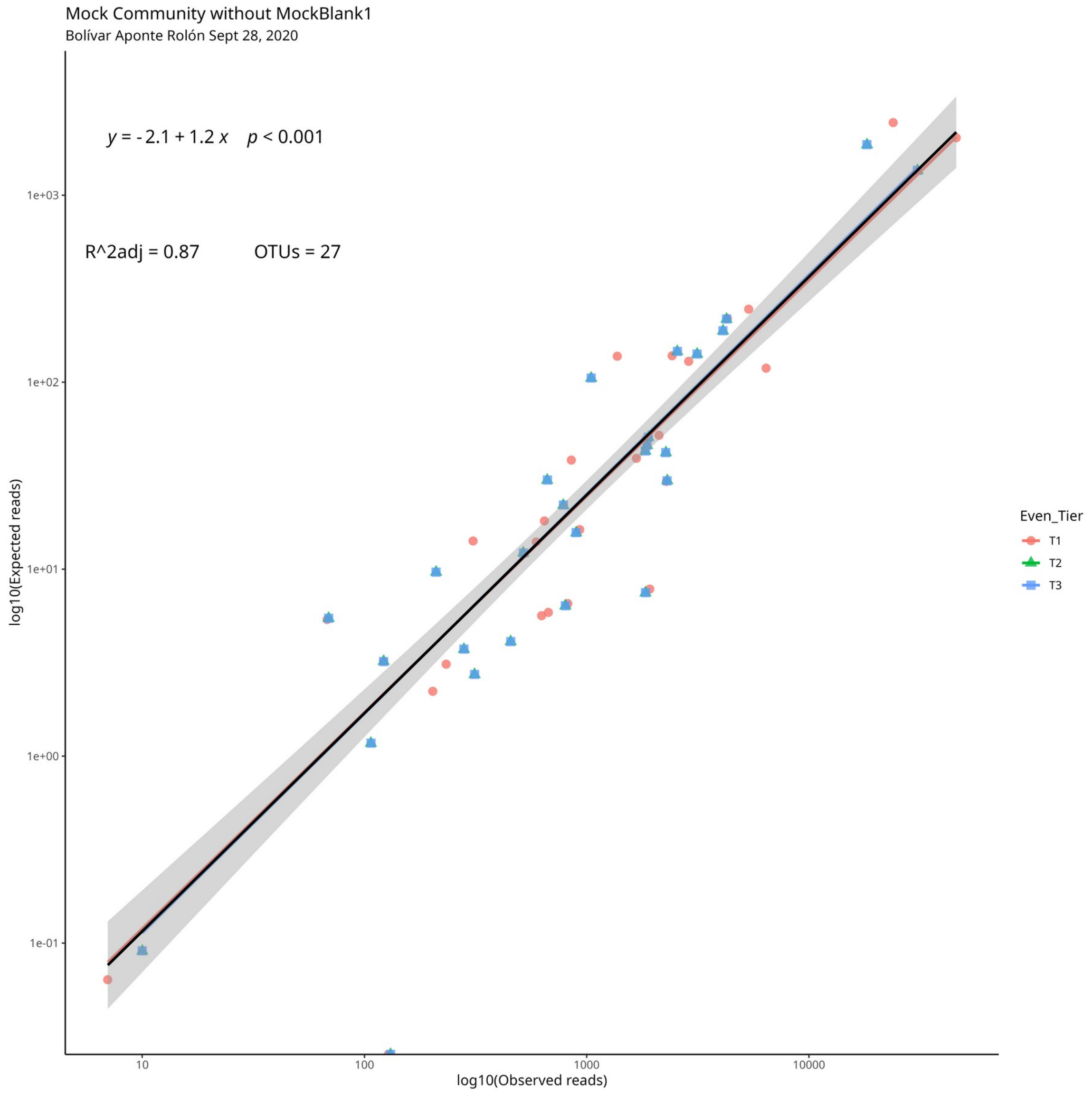


*Simple linear regression of tiered mock community.*

#### Supplementary Material - Methods

*Seed collection and sterilization*

Seeds were prepared for germination within 24 hours of fruit collection on BCI, January - April 2019. Fruits were peeled to extract seeds, cleaned, put in water to swell embryo, sterilized, and planted in sterile soil germination trays (75% soil; 25% sand). Soil was autoclaved at 121 °C in two, one hour cycles. Seeds were surface sterilized using 10% bleach for 3 minutes followed by 70% ethanol for 3 minutes.

*Germination Tray and 24-pot tray sterilization*

Plastic germination trays and 24-cell trays were sterilized in a 10% bleach bath for 20 minutes, sprayed with 70% ethanol, and paper towel dried right before adding soil/planting.

*Planting in germination trays*

The sterile soil is added to an sterilized plastic germination tray, combined with water well until it is wet and has a cookie dough-like consistency. Seeds were then added to moist soil. A sprinkle of dry soil is added to as a top layer to discourage pathogen spores from landing in the wet surface soil.

*19.0.4 Seedling transfer into pots*

Once seeds germinated, seedlings were transferred to pots by wetting the soil and extracting intact root system. Seedling were immediately placed in pot with sterile soil using a small shovel after uprooting from germination tray. Hands were sprayed with 70% ethanol when switching from handling one species to handling another. Plants were watered as needed at the soil level and 25mL of MiracleGro all-purpose plant food was added to every seedling once a month throughout entirety of experiment.
